# VISTA: An integrated framework for structural variant discovery

**DOI:** 10.1101/2023.08.11.553053

**Authors:** Varuni Sarwal, Seungmo Lee, Jianzhi Yang, Sriram Sankararaman, Mark Chaisson, Eleazar Eskin, Serghei Mangul

**Author notes:** Denotes equal contribution (Varuni Sarwal, Seungmo Lee). Denotes joint supervision (Eleazar Eskin, Serghei Mangul).

## Abstract

Structural variation (SV), refers to insertions, deletions, inversions, and duplications in human genomes. With advances in whole genome sequencing (WGS) technologies, a plethora of SV detection methods have been developed. However, dissecting SVs from WGS data remains a challenge, with the majority of SV detection methods prone to a high false-positive rate, and no existing method able to precisely detect a full range of SV’s present in a sample. Previous studies have shown that none of the existing SV callers can maintain high accuracy across various SV lengths and genomic coverages. Here, we report an integrated structural variant calling framework, VISTA (Variant Identification and Structural Variant Analysis) that leverages the results of individual callers using a novel and robust filtering and merging algorithm. In contrast to existing consensus-based tools which ignore the length and coverage, VISTA overcomes this limitation by executing various combinations of top-performing callers based on variant length and genomic coverage to generate SV events with high accuracy. We evaluated the performance of VISTA on using comprehensive gold-standard datasets across varying organisms and coverage. We benchmarked VISTA using the Genome-in-a-Bottle (GIAB) gold standard SV set, haplotype-resolved de novo assemblies from The Human Pangenome Reference Consortium (HPRC)^1,2^, along with an in-house PCR-validated mouse gold standard set. VISTA maintained the highest F1 score among top consensus-based tools measured using a comprehensive gold standard across both mouse and human genomes. VISTA also has an optimized mode, where the calls can be optimized for precision or recall. VISTA-optimized is able to attain 100% precision and the highest sensitivity among other variant callers. In conclusion, VISTA represents a significant advancement in structural variant calling, offering a robust and accurate framework that outperforms existing consensus-based tools and sets a new standard for SV detection in genomic research.

## Introduction

Structural variants (SVs) are genomic regions that contain an altered DNA sequence due to deletion, duplication, insertion, inversions and other complex rearrangements. SVs are present in approximately 1.5% of the human genome^3^, but this small subset of genetic variation has been implicated in the pathogenesis of psoriasis^4^, Crohn’s disease^5^ and other autoimmune disorders^6^, autism spectrum and other neurodevelopmental disorders^7–10^, and schizophrenia^11–14^. Specialized computational methods—often referred to as SV callers—are capable of detecting structural variants directly from sequencing data. Although a plethora of SV detection methods have been developed, dissecting SVs from WGS data presents a number of challenges, with the majority of SV detection methods prone to from a high false-positive rate^12,13^, and no existing method able to accurately detect a full range of SV’s present in a sample.

To address these challenges, we developed VISTA (Variant Identification and Structural Variant Analysis), an automated SV detection algorithm integrating multiple variant callers. VISTA automatically executes the combination of individual callers and is able to achieve a balance between sensitivity and precision. We have extensively the performance of VISTA using three comprehensive datasets with high accurate gold standard benchmarking data across 17 WGS samples, namely the Genome-in-a-Bottle (GIAB) gold standard SV set, haplotype-resolved de novo assemblies from The Human Pangenome Reference Consortium^1,2^ (HPRC), along with an in house PCR validated gold standard across seven mouse strains to demonstrate its superiority, in consistently maintaining the highest F1 score among top consensus-based tool. Notably, HPRC represents a highly accurate SV set consisting of haplotype-resolved de novo assemblies and was never used for a comprehensive assessment of the accuracy of SV callers. Using prepared benchmarks, we compare the performance of VISTA with 21 individual SV callers (Octopus^15^, Pindel^16^, Manta^17^, CLEVER^18^, DELLY^19^, PopDel^20^, BreakDancer^21^, GASV^22^, Smoove^23^, GenomeSTRiP^24^, MiStrVar^25^, indelMINER^26^, GRIDSS^27^, Tardis^28^, CREST^29^, RDXplorer^30^, transIndel^31^, LUMPY^32^, GROM^33^), and 3 popular consensus-based callers (Parliament2^34^, SURVIVOR^35^ and Jasmine^36^) to demonstrate VISTA’s ability to balance precision and sensitivity with the highest F-score, consistently across different organisms, variant length categories and data of varying genomic coverages. VISTA obtains the highest F-score of 0.785 on human full coverage data, of 0.769 on full coverage mouse data and of 0.51 on WGS data with coverage less than 0.5x. VISTA represents a transformative step forward in the field of structural variant calling, providing a powerful and accurate framework that surpasses existing consensus-based tools and establishes a new benchmark for SV detection in genomic research.

## Results

### Existing methods for structural variant calling

Detecting SVs using short read sequencing is a challenging problem, and a variety of different SV detection methods have been developed. Typically, SVs are detected by looking for changes in read depth (RD) (eg: GROM^33^), identifying clusters of discordantly aligned paired-end (PE) reads (BreakDancer^21^) or split reads (SRs) (eg: Octopus^15^), constructing assemblies or a combination of these approaches. Read-pair methods leverage information about deviant pair distances and orientations in paired-end sequencing data. Read-depth methods are based on the assumption that the sequencing depth over a genomic region is proportional to its copy number. Consequently, deviations from the expected depth may indicate a deletion or duplication. Split-read methods utilize reads that are partially aligned to the reference genome, indicating a potential structural variant at the breakpoint. Assembly-based methods create a de novo assembly of the genome and then align the assembly to a reference genome to detect structural variants. Hybrid methods combine one or more of the above approaches to increase sensitivity and specificity. This diversity of approaches also results in performance heterogeneity across SV types and sizes, as well as varied compute requirements. Consensus-based SV callers, such as Parliament2^34^, Jasmine^36^, and SURVIVOR^35^ callers perform ensemble optimization by combining the outputs of multiple SV callers into a high-quality consensus set.

### VISTA: An integrated framework for structural variant discovery

VISTA is a novel consensus-based integrated framework for structural variant discovery, designed to overcome the challenges associated with variant calling. Leveraging multiple variant callers based on organism characteristics, genomic coverage, and variant length, VISTA dynamically adapts to diverse genomic contexts, ensuring precise and comprehensive SV detection. VISTA’s unique merging procedure combines the outputs of individual callers using a novel filtering and merging algorithm, resulting in a highly accurate SV set with minimized false positives and false negatives. Additionally, VISTA offers an optimized mode for prioritizing either precision or recall, further enhancing its flexibility and applicability to diverse research objectives.

### Datasets used for this study

In order to assess the accuracy of VISTA, we used datasets obtained from various organisms, read lengths, and genomic coverages. We used 17 WGS samples from three major datasets, referred to as D1, D2, and D3. These datasets were used to compare the performance of VISTA with 18 individual SV callers and 3 consensus-based callers, using comprehensive gold standards, namely Genome in a Bottle (GIAB-HC), Human Pangenome Reference Consortium (HPRC-HC) and Mus musculus (MM-7). D1 consists of the H002 Ashkenazi Jewish Trio son sample from the Genome-in-a-Bottle (GIAB) consortium^23^. The alignment files were publicly available from the GIAB website and were used as input to the SV callers. The average depth of coverage was 45x and the reads were 2×250 bp. For GIAB-HC, we use the GIAB preliminary variant set containing 37412 deletions in HG002. We then extracted the high confidence regions using the high confidence bed file and variants greater than 50 bp, to obtain our gold standard set of deletions consisting of 4120 deletions (GIAB-HC-D). For insertions, we used the GIAB preliminary variant set consisting of 35163 insertions (GIAB-I). D2 refers to ten WGS human samples from the 1000 Genomes Project. The average depth of coverage was 30x and the reads were 2×150 bp. For the HPRC-HC dataset, we used haplotype-resolved de novo assemblies from the Human Pangenome Reference Consortium (HPRC) to generate benchmarking SV sets. On average, each set contains 17807 deletions larger than 50 bp (Supplemental Figure 4(c)). Dataset D3 refers to seven inbred mouse WGS samples. The high coverage sequence data was used as an input to the SV callers in the form of aligned reads for all experiments. For 7MM-HC^37^ we used a PCR-validated set of deletions, in which the mouse deletions were manually curated, and targeted PCR amplification of the breakpoints and sequencing was used to resolve the ends of each deletion to the base pair (Supplemental Table 2).

### Evaluation metrics

Callers were evaluated based on their ability to detect structural variants using precision, sensitivity and f-score as metrics. These metrics were computed using a system of resolution thresholds that varied from 10 to 10000 bp. An inferred deletion was considered to be correctly predicted if the distance of right and left coordinates is within the resolution threshold from the coordinates of true deletion.

### VISTA achieves the optimal balance of precision and sensitivity for deletions calls across two independent human benchmarks

We assessed the performance of VISTA in terms of precision (false discovery rate), recall (true-positive rate), compared to other individual (eg DELLY^19^, Manta^17^, Lumpy^32^) and consensus-based (eg Parliament2^34^, SURVIVOR^35^ and Jasmine^36^) short-read SV methods across 11 WGS samples from D1-D2 datasets with gold standard sets available as a part of GIB-HC-D and HPRC-HC.

First, we compared the length distribution of the deletions in the GIB-HC, compared to VISTA and other callers. VISTA and Manta^17^ were the top 2 callers with a median length closest to the gold standard (Figure 1a). Notably, all of the consensus-based variant callers overestimate the deletion length by 175 bp on average. Parliament2^34^ was the only consensus caller that overestimated the number of deletions, while all other callers underestimated. We plotted Venn diagrams to better visualize overlapping deletions call sets for VISTA, the gold standard, and other top-performing variant callers for HG002 chr19 (Figure 1 c-e). Callers either had too many false positives (Parliament2^22^, Jasmine^24^), detected too few deletions (SURVIVOR^23^), or had a smaller overlap with the ground truth (Manta^5^). We observed a high concordance of deletions predicted by VISTA and the ground truth set. Next, we compared the performance of VISTA with other SV callers in terms of inferring deletions for a range of resolution thresholds from 10 to 10000 bp (Figure 1f-h). VISTA has achieved the highest recall at 100bp (72.4%) while having the fourth-highest precision (85.7%). Only SURVIVOR^35^ (95.9%), Manta^17^ (93.4%) and Tardis^38^ (86.3%) had a higher precision, at the cost of a lower recall: SURVIVOR^35^ (48.3%), Manta^17^ (61.0%), Tardis^28^ (16.8%). Importantly, VISTA achieves the highest F1 score (78.5%) for thresholds 100bp and above, followed by Manta^17^(73.8%) and Jasmine^36^ (72.4%) (Figure 1h). For a threshold of 10bp, Manta^17^(69.5%) had a marginally higher F-score than VISTA (63.8%), followed by Jasmine^36^(63.6%).

**Figure 1:**
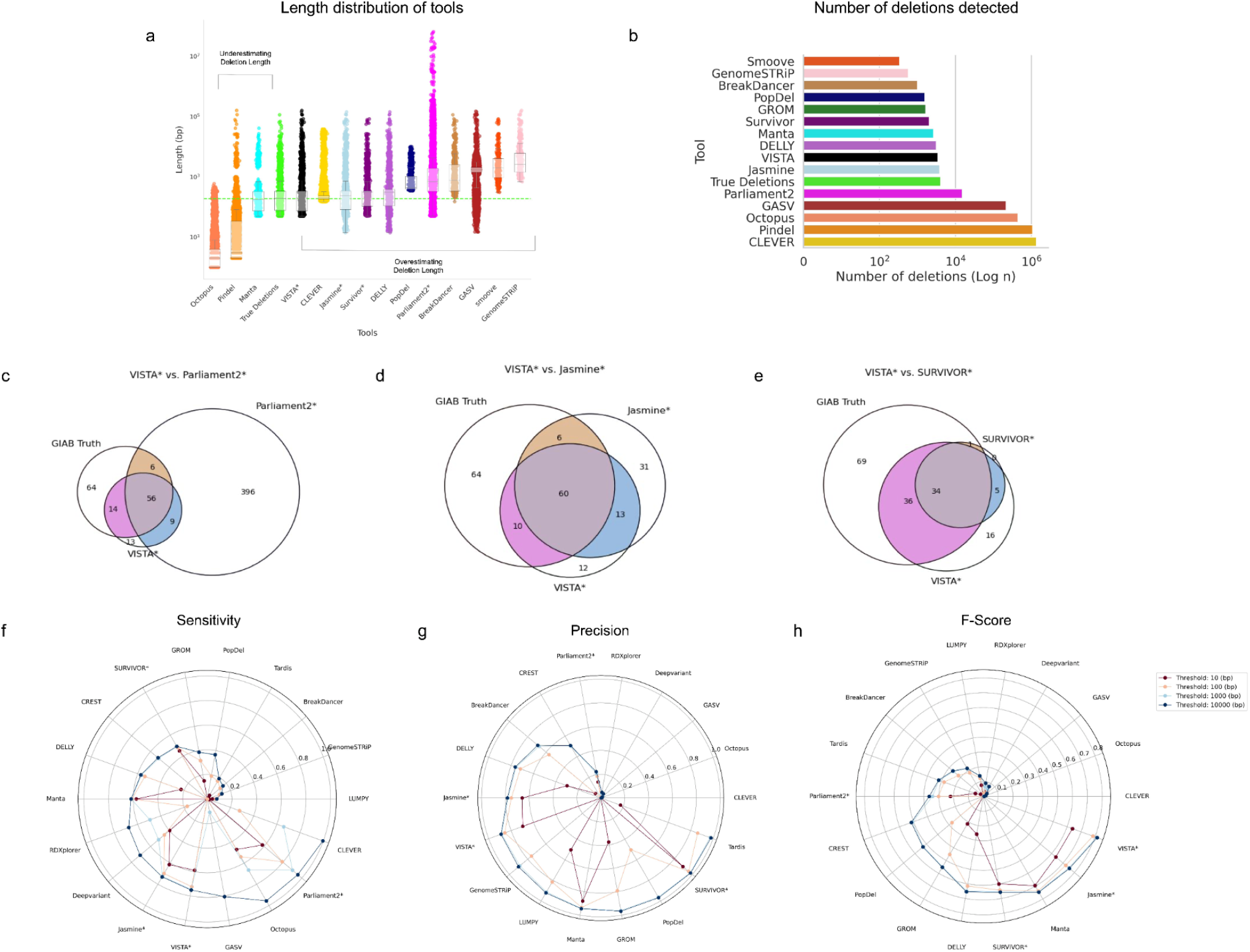
Comparing the performance of VISTA with popular SV callers on GIAB-HC-D. A deletion is considered to be correctly predicted if the distance of right and left coordinates are within the threshold from the coordinates of a true deletion (a) Length distribution of VISTA vs GIAB-HC-D and other callers (b)Number of deletions detected by VISTA vs GIAB-HC-D and other callers (c-e) Venn diagrams of intersecting top SV call sets on HG002 chr 19. (f) Sensitivity of SV callers at different thresholds. (g) Precision of SV callers at different thresholds. (h) F-score of SV callers at different thresholds. The asterisk represents consensus-based callers.

Next, we studied the robustness of VISTA across 10 human samples on the HPRC dataset (Figure 2). VISTA had the closest median deletion length compared to the gold standard (Figure 2a). All of the consensus based callers underestimated the number of deletions (Figure 2b). We observed all the callers to have a consistent trend across the samples, with VISTA consistently achieving the highest F-score across all samples (Figure 2e). Most samples demonstrated a slightly elevated precision for samples HG00735 and HG01234, and a minima at sample HG01175 (Figure 2d).

**Figure 2:**
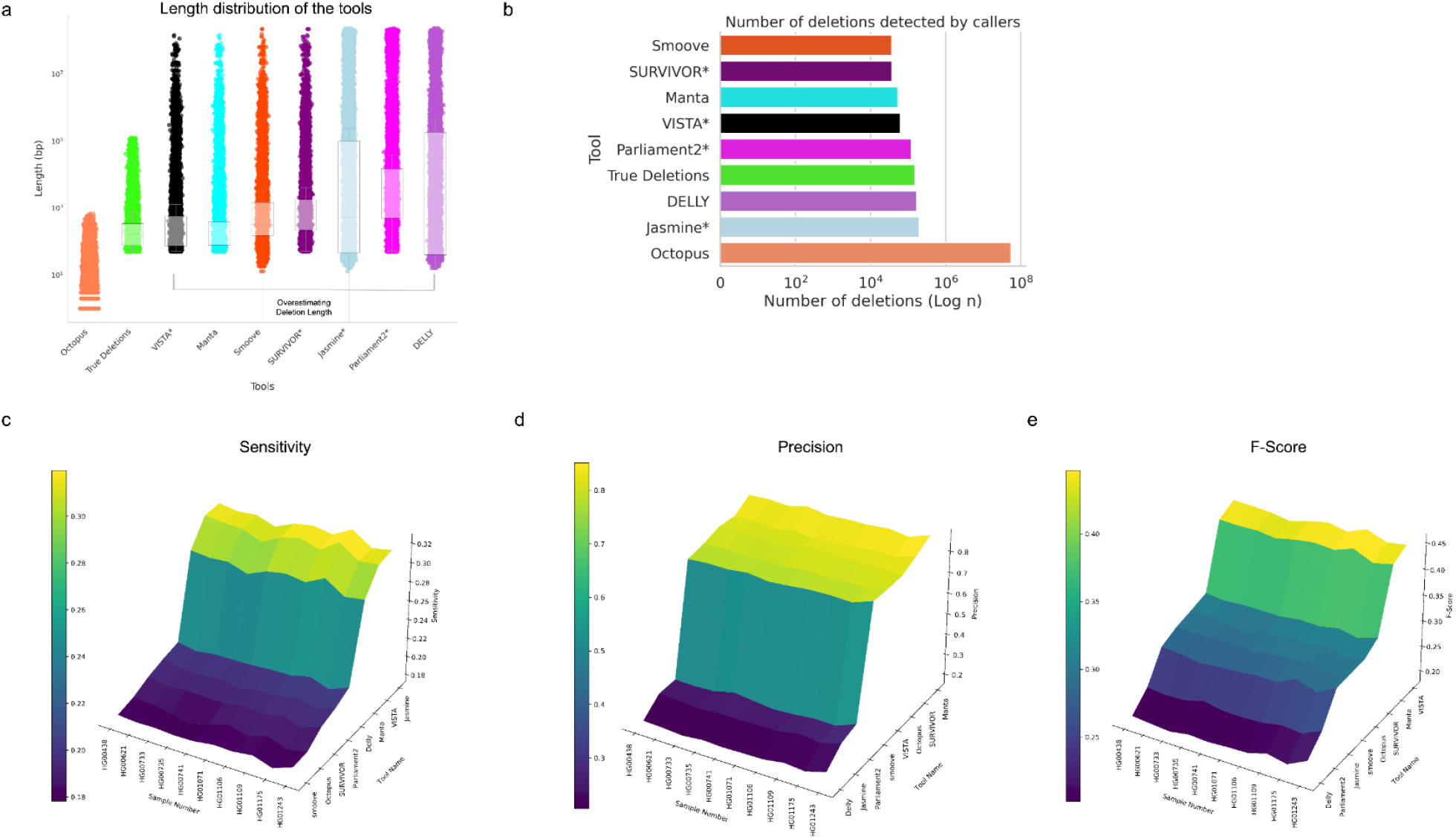
Comparing the performance of VISTA with popular SV callers on HPRC-HC. A deletion is considered to be correctly predicted if the distance of right and left coordinates are within the threshold from the coordinates of a true deletion (a) Length distribution of VISTA vs HPRC-HC and other callers (b)Number of deletions detected by VISTA vs HPRC-HC and other callers (f) Sensitivity of VISTA vs other callers for 10 samples (g) Precision of VISTA vs other callers for 10 samples (h) F-score of VISTA vs other callers for 10 samples. The asterisk represents consensus-based callers.

### Vista maintains high performance for deletions calls across Mus musculus samples

We compared the performance of VISTA with 16 individual SV callers and 3 consensus-based SV callers in terms of inferring deletions on 7MM-HC. We compared the deletion lengths predicted by VISTA and other consensus callers to the ground truth. VISTA was the closest in terms of the length distribution of deletions as compared to 7MM-HC (Figure 3a). We plotted Venn diagrams to better visualize overlapping deletions call sets for VISTA, the gold standard, and other top-performing variant callers for 7MM chr19 (Figure 3 c-e). Similar to human data, callers either had too many false positives (Parliament2^22^, Jasmine^24^), detected too few deletions (SURVIVOR^23^). We observed a high concordance of deletions predicted by VISTA and the ground truth set. We analyzed the performance to detect mouse deletions for a range of resolution thresholds from 10 to 10000 bp (Figure 3 f-h). VISTA has the third highest precision (77%) at 100bp and above (Figure 3g). Only TARDIS^28^ (78%) and PopDel^20^ (79%) are marginally higher, at the cost of a significantly lower recall. Several tools such as Jasmine^36^(70%), GRIDSS^27^ (63%) outperform VISTA (61%) in terms of Sensitivity but are among the lowest performing tools for Precision (Figure 3f). Thus, VISTA (68%) has the highest F1 score for thresholds 100bp and above, followed by Manta^17^ (65.0%) and LUMPY^32^ (64.8%) (Figure 3h). For a threshold of 10bp, LUMPY^32^ (19%), SURVIVOR^35^ (17%), and Manta^17^ (11.2%) have a marginally higher F-score than VISTA (10.6%). Among the consensus-based callers, SURVIVOR^35^ achieves the highest F-score (63%) at 100 bp and above, followed by Parliament2^34^ (48%) and Jasmine^36^ (44%). Results across the individual mouse strains are depicted in Supplemental Figure 2.

**Figure 3:**
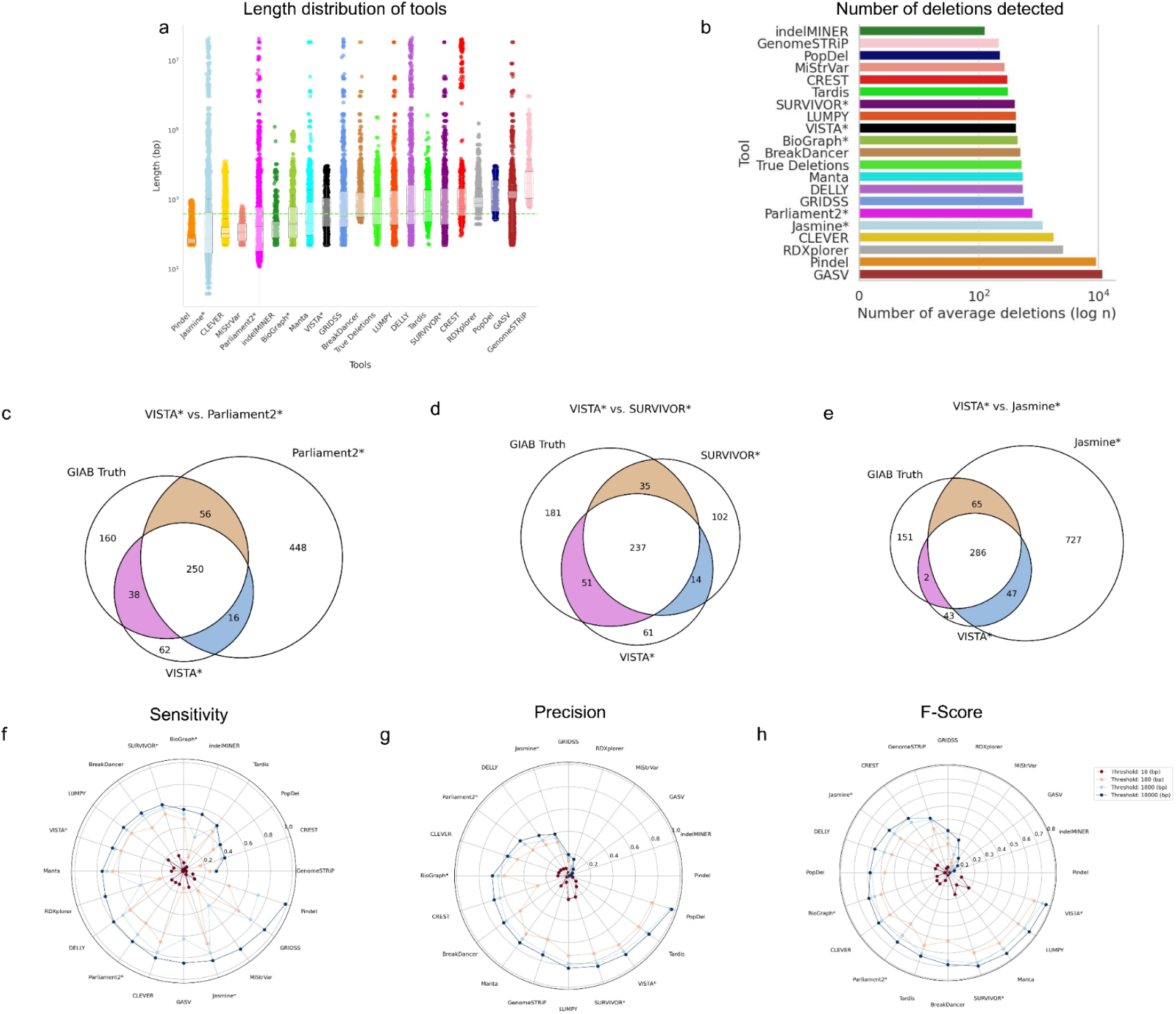
Comparing the performance of VISTA with popular SV callers on mouse data. A deletion is considered to be correctly predicted if the distance of right and left coordinates are within the threshold from the coordinates of a true deletion. (a) Sensitivity of SV callers at different thresholds. (b) Precision of SV callers at different thresholds. (c) F-score of SV callers at different thresholds. Results displayed above report the average across 7 different mouse strains. d) Venn diagrams of GIAB truth and VISTA with other top callers on AKR_J mouse strain. The asterisk represents consensus-based callers.

### Vista delivers highly accurate insertion calls

We compared the performance of VISTA with several other consensus-based callers in terms of inferring insertions on GIAB-I. In contrast to deletions, the length distribution of detected insertions varied across tools, and was substantially different from the true distribution (Figure 4a). Notably, none of the callers demonstrated the capability to accurately predict long insertions (>1000bp). Hence, all the callers underestimated the number of insertions (Figure 4b). We found VISTA and Parliament2^34^ to be the two callers with median insertion lengths of 16 bp and 66 bp closest to the gold standard (44 bp) (Figure 4a). To assess the performance of VISTA against other callers in detecting insertions, we analyzed a range of resolution thresholds spanning from 10 to 10000 bp (Figure 4c-e). VISTA has the highest recall at 100bp (6.3%) while having the fifth-highest precision (65%) at 100bp. While both TransIndel^31^ (95.40%) and Manta^17^ (94.31%) had a higher precision, this came at the expense of a lower recall. Thus, VISTA (11.6%) had the highest F1 score for thresholds 10 bp and above, followed by Parliament2^34^ and Manta^17^(9.8%) at 100 bp.

**Figure 4:**
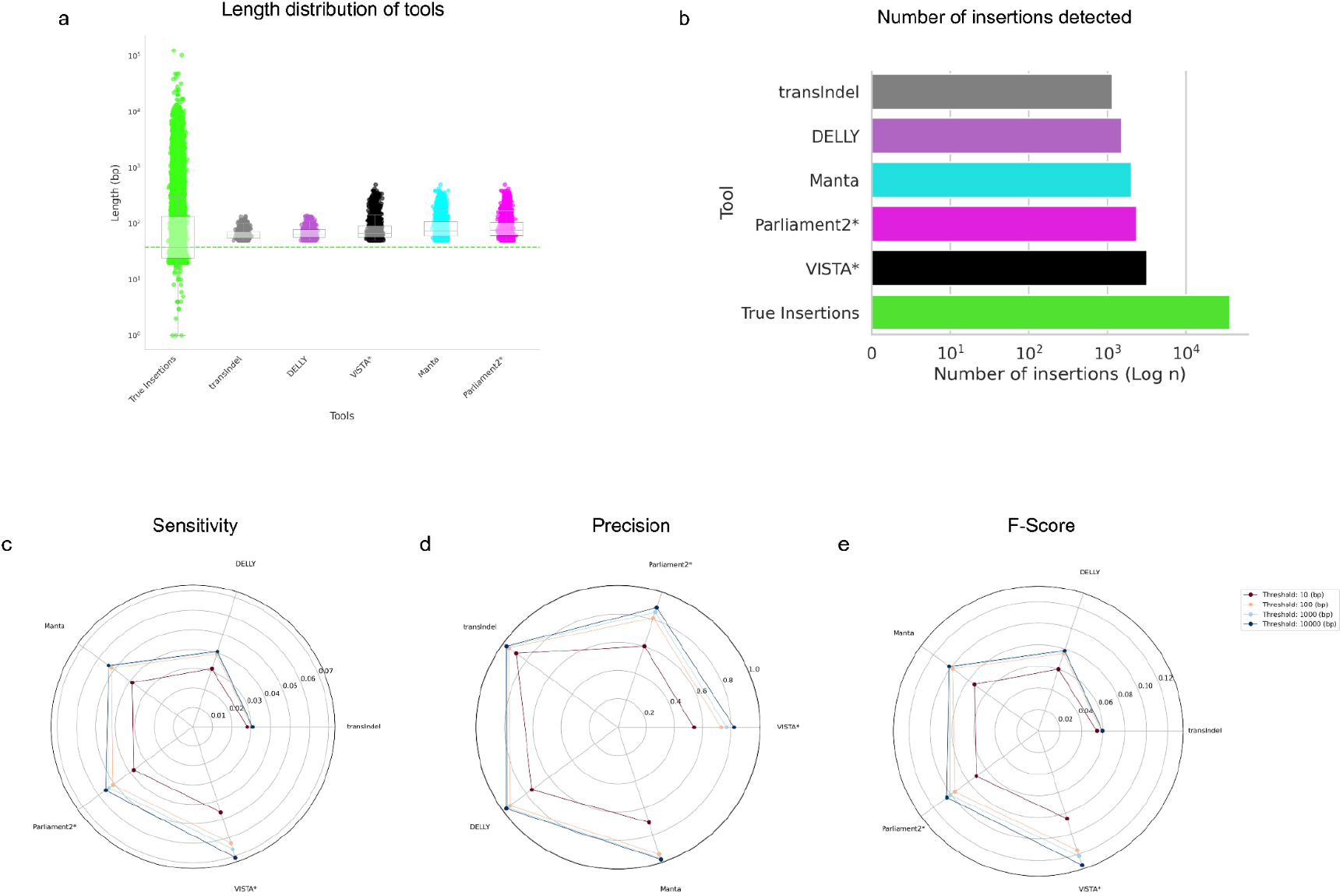
Comparing the performance of VISTA with popular SV callers on GIAB-HC-I. An insertion is considered to be correctly predicted if the distance of right and left coordinates are within the threshold from the coordinates of a true insertion (a) Length distribution of VISTA vs GIAB-HC-I and other callers (b)Number of insertions detected by VISTA vs GIAB-HC-I and other callers (c) Sensitivity of SV callers at different thresholds (d) Precision of SV callers at different thresholds (e) F-score of SV callers at different thresholds. The asterisk represents consensus-based callers.

### Vista achieves the highest performance on low coverage WGS samples

We assessed the performance of VISTA as well as other SV callers at different coverage depths generated by down-sampling the original mouse WGS data (Figure 5). The simulated coverage ranged from 32x to 0.1x, and ten subsamples were generated for each coverage range. For each method, the number of correctly detected deletions generally decreased as the coverage depth decreased. VISTA was able to obtain the highest F-score consistently across all coverages from 0.5x to 32x, closely followed by Delly for 0.5x, SURVIVOR^35^ and Jasmime^36^ for 1x-8x. For higher coverages ranging from 16x-32x, SURVIVOR^35^ and Manta^17^, and LUMPY^32^ were the top 3 tools closest in performance to VISTA (Figure 5c). VISTA obtained a maximum precision of 0.73 at an intermediate coverage of 4x (Figure 5b). While methods like LUMPY^32^ were able to achieve a higher precision of up to 0.94, this came at a cost of a lower sensitivity. As the coverage increased, the sensitivity and F-score increased, with a maximum value of 0.64 and 0.66 respectively, at 32x coverage (Figure 5a).

**Figure 5:**
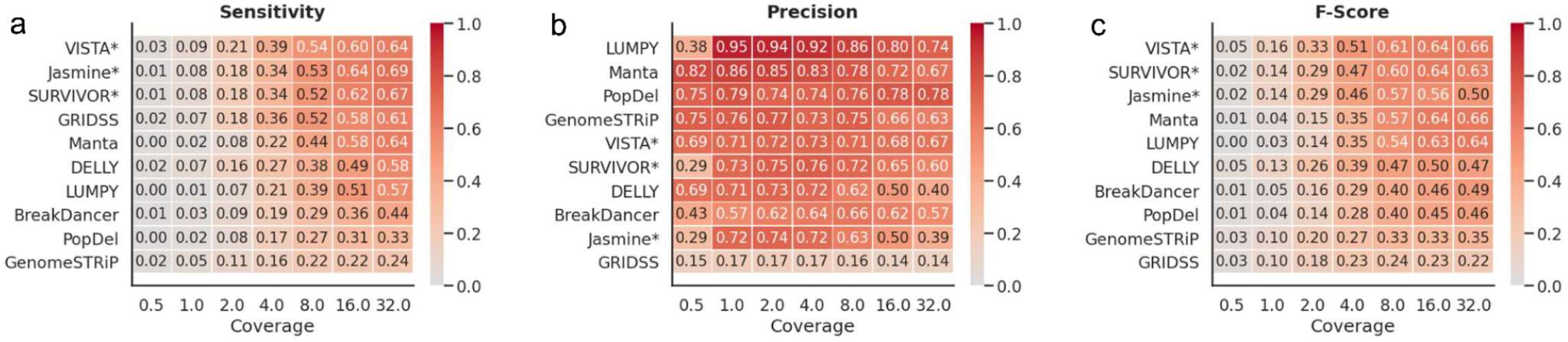
Performance of VISTA on low and ultra-low coverage mouse data. (a) Heatmap depicting the sensitivity based on 100 bp threshold across various levels of coverage. (b) Heatmap depicting the precision based on 100 bp threshold across various levels of coverage. (c) Heatmap depicting the F-score based on 100 bp threshold across various levels of coverage. The asterisk represents consensus-based callers.

### VISTA offers an additional mode to boost previsions or sensitivity

We recognize that certain applications may require a highly precise variant caller or a caller with a high recall. Hence, we have provided an “optimize mode” in VISTA, where the user can choose to optimize for either precision or recall. We compared the sensitivity of VISTA-optimized as compared to other SV callers (Figure 6a). VISTA (85%) obtains the highest sensitivity consistently across all thresholds from 10 to 10000 bp and is significantly higher than the second and third top-performing callers, Jasmine^36^ (70%) and Manta^17^ (65%). We also compared the precision of VISTA compared to other SV callers (Figure 6b), and VISTA (100%) obtains the highest precision at 100 bp and above with no false positives, closely followed by SURVIVOR^35^ (95%) and Manta^17^(93%).

**Figure 6:**
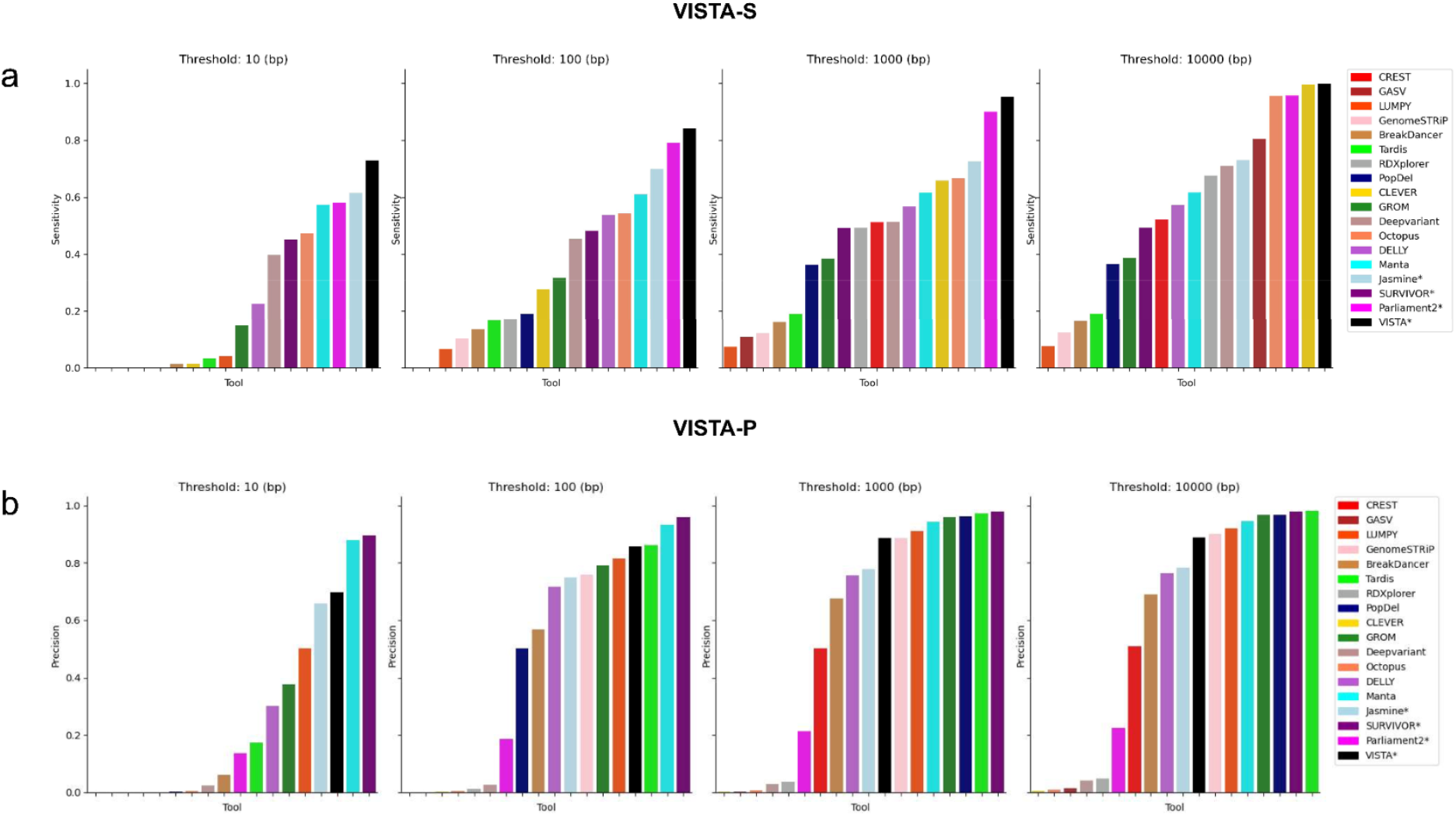
Comparing the performance of VISTA-optimized with popular SV callers on human data. A deletion is considered to be correctly predicted if the distance of right and left coordinates are within the threshold from the coordinates of a true insertion. (a) Sensitivity of VISTA-S vs other SV callers at different thresholds. (b) Precision of VISTA-P vs other SV callers at different thresholds

### Computational performance of VISTA

We compared the central processing unit (CPU) and random access memory (RAM) requirements of VISTA with other popular consensus-based callers (Figure 10). VISTA and SURVIVOR^35^ were the two callers with the lowest RAM usage, with 7 MB of RAM used to process the human sample from GIAB-HC-D dataset. Additionally, VISTA has a significantly low CPU time, of 0.65 seconds, as compared to Parliament2^34^ (41.7 hours) and Jasmine^36^ (11.5 seconds). We acknowledge that certain consensus callers such as VISTA, Jasmine and SURVIVOR take individual callers and input and only account for the consensus algorithm in the run time. On the other hand, callers like Parliament2 run the individual callers through a docker based image, resulting in a higher run time. We observed VISTA to be highly scalable across 10 samples, data of varying coverage.

## Discussion

Here, we report the development and evaluation of VISTA (Variant Identification and Structural Variant Analysis), an integrated structural variant calling framework, that leverages the results of individual callers using a novel and robust filtering and merging algorithm. Our study is the first to comprehensively benchmark the performance of a novel SV detection method against the HPRC benchmark. We demonstrate VISTA’s abilities in being the only SV caller that is able to obtain a consistently high performance across variant types, datasets, coverages, and organisms.

While the recall ability for insertions is reduced due to limitations of short-read–based insertion detection algorithms, VISTA was able to obtain the highest F-score among other popular individual and consensus-based callers. We attempted to additionally train the insertion call set on SVseq232 and inGAP-sv33, but these tools fail to detect both the start and end positions of insertions. All the SV callers we used for training had limitations on finding large insertions, where the largest insertion size Manta outputs predicted is 503 bp, and the gold standard has 3568 insertions that exceed 500 bp across the full chromosome. Ultimately, VISTA merges Manta4 and transIndel19 to maximize the F-score and outperform other callers.

We observed Parliament2’s output to be highly inconsistent across human samples. For certain samples, it was only able to run a subset of the callers (Supplemental Table 6). DELLY^6^, CNVnator^27^, and LUMPY^20^ were the only 3 callers that Parliament2 ran across all 10 human samples. In order to provide a fair comparison across the samples, we only considered the outputs of the three callers. While we used a custom script to compare the performance of the callers with the gold standard, we reran our analysis using Truvari^28^, a tool widely adopted by the community for the evaluation of SV calls, and found our results to be consistent. In this study, we did not filter the output of the SV callers. Filtering is extremely context-specific and may vary significantly for different experiments, and good post-hoc filtering would require individual consideration of each tool’s quality metrics and thresholds to get comparable results.

While VISTA demonstrates robust performance when applied to datasets it was pre-trained on, we observed limitations in its ability to generalize well to new datasets not included in the training data. For instance, when VISTA was trained on the HPRC dataset, it achieved the highest performance when evaluated on 10 different human samples with comparable characteristics (Figure 2). However, when VISTA pre-trained on the HG002 sample was applied to the HPRC dataset, which it was not pre-trained on, the results were not as optimal (Supplemental Figure 6). This suggests that pre-training plays a major role in VISTA’s, and caution should be exercised when applying it to novel datasets with significantly different characteristics.

**Table 1:** Datasets used in this study grouped by the number of samples and sample names (indicated in the columns “# Samples” and “WGS Samples”. We documented how each of the gold standard datasets were prepared (“Method of preparation”), and recorded the read length and organism corresponding to the dataset. We also recorded the paper where the dataset was first published (“Citation”).

| Dataset name | # Samples | Method of preparation | Organism | WGS samples | Reference | Read length |
| --- | --- | --- | --- | --- | --- | --- |
| GIAB high confidence set (GIAB-HC) | 1 | integrate multiple short- and linked-read sequencing datasets | Human | HG002 | Zook, Justin M., et al. "An open resource for accurately benchmarking small variant and reference calls." *Nature biotechnology* 37.5 (2019): 561-566. | 2x250 bp |
| Human Pangenome Reference Consortium (HPRC-HC) | 10 | Haplotype-resolved de novo assemblies from HPRC | Human | HG00733, HG00438, HG00621, HG00735, HG00741, HG01071, HG01106, HG01109, HG01243, HG01175 | Wang, Ting, et al. "The Human Pangenome Project: a global resource to map genomic diversity." *Nature* 604.7906 (2022): 437-446. | 2x150bp |
| Mouse gold standard set (7MM-HC) | 7 | In-house PCR validated | Mouse | A_J, AKR_J, BALB_CJ, C3H_HeJ, CBA_J, DBA_2J and LP_J. | Sarwal, Varuni, et al. "A comprehensive benchmarking of WGS-based deletion structural variant callers." *Briefings in Bioinformatics* 23.4 (2022): bbac221. | 2x100 bp |

## Methods

### Preparing the GIAB-HC dataset

We used public benchmark data for the Ashkenazi Jewish Trio son (NA24385/HG002) from the Genome-in-a-Bottle (GIAB) consortium. The GIAB-HC SV truth set is based on HG002, a male Ashkenazi Jewish sample using multiple technologies and manual vetting of the SV. While VISTA can infer multiple SV types, the current GIAB call set largely comprises insertion and deletion events. The alignment files were publicly available from the GIAB website and were used as input to VISTA and other variant callers. The average depth of coverage was 45x and the reads were 2×250 bp paired-end reads. We used the Genome-in-a-Bottle preliminary variant set containing variants in HG002 as our gold standard. The preliminary HG002 variant set available is the first reference set that is near complete within defined high-confidence regions of the genome defined by a bed file and hence allowed us to systematically benchmark the performance of variant callers within those high confidence regions. The set contained 71152 variants, out of which 8190 deletions remained after extracting variants with length over 50 bp, and 4120 deletions remained after filtering the high-confidence regions. Out of 4120 filtered deletions, 44.5% of the deletions were in the range of 100-500 bp. 11.5% of deletions were larger than 1000 bp (Figure 6). Furthermore, 14779 insertions remained after filtering insertions with lengths greater than 50 bp, and 6723 insertions remained after filtering the high-confidence regions. Complete details on how the human gold standard was prepared are provided in the Methods section.

### Preparing HPRC-HC dataset

In order to demonstrate the scalability of VISTA across a large number of samples and gold standard sets, we tested VISTA on the following samples of human data: HG00733, HG00438, HG00621, HG00735, HG00741, HG01071, HG01106, HG01109, HG01243, HG01175. The alignment files are from the 1000 Genomes Project with 30x coverage and 2×150bp reads on the reference GRCh38^39^. They are publicly available through The International Genome Sample Resource (IGSR) (Fairley et al., 2020) portal. VISTA and other variant callers take these alignment bam files as input and output SV sets.

Since haplotype-resolved assemblies can span and capture longer SVs especially in repetitive and complex regions, they have been used to create benchmark SV sets^40^. The recent production of high-quality assemblies provide accurate information of the sample genomes. We used dipcall^41^ to generate SVs of the human samples directly from haplotype-resolved de novo assemblies produced by The Human Pangenome Reference Consortium (HPRC). On average, each dipcall sample SV set contains 588783 deletions and 17807 deletions are larger than 50 bp. 8165 deletions are in the range of 100-500 bp and 1948 deletions are larger than 1000 bp. These SVs from dipcall were used as a benchmark to test VISTA on the 10 human samples.

### Preparing the 7MM-HC dataset

In order to demonstrate VISTA’s generalizability across organisms, we used a set of homozygous deletions present in inbred mouse chromosomes. More specifically, we chose to limit our analysis to mouse chr19 since it is the smallest. We used a PCR-validated set of deletions, in which the mouse deletions were manually curated, and targeted PCR amplification of the breakpoints and sequencing was used to resolve the ends of each deletion to the base pair. The same read alignment file which was used for manual curation of the deletions was used as an input to the structural variant callers, making our gold standard complete, containing all possible true deletions (true positives). To ensure that our gold standard is complete with respect to the alignment file, we first examined all possible deletions manually and then validated each deletion by PCR. Thus, while our gold standard may not be universally complete, our gold standard was complete with respect to the alignment files which were provided to the SV callers since all deletions which could be possibly detected from the alignment were recorded and further examined using PCR. While homozygous strains do not fully reflect the biology of real data, we use it in our mouse model as an easy-to-detect simple baseline. The set of deletions we used among seven inbred strains, called with reference to C57BL/6J, is illustrated in Figure 1a and listed in Supplemental Table 5. We filtered out deletions shorter than 50 bp, as such genomic events are usually detected by indel callers rather than SV callers. In total, we obtained 3,710 deletions with lengths ranging from 50 to 239,572 base pairs (Supplemental Figure 3 and Supplemental Table 5). Almost half of the deletions were in the range of 100-500 bp. 27.9% of deletions were larger than 1000 bp (Supplemental Figure 4a). High coverage sequence data was used as an input to the SV callers in the form of aligned reads. Reads were mapped to the mouse genome (GRCm38 Mouse Build) using BWA with -a option. In total, we obtained 5.2 billion 2×100 bp paired end reads across seven mouse strains. The average depth of coverage was 50.75x. Details regarding the gold standard and raw data preparation and analysis are presented in the Supplementary Materials.

### VISTA algorithm

VISTA is a consensus-based variant caller that produces high accuracy call sets based on the input of individual variant callers using a novel merging and filtering algorithm (Supplemental Figure 1). VISTA takes the output of individual callers as inputs, and produces a highly accurate file predicting variants, in vcf format. VISTA consists of two states, pre-training and discovery. In the pre-training stage, we use datasets with a well defined gold standard. First, a list of input vcf files are provided to VISTA as input, containing variants called by individual top-performing callers. Next, VISTA identifies which organism is being called, the genomic coverage of the input sample, as well as the length distribution of the predicted variants. Then, based on the parameters above, namely the organism type, coverage and length, VISTA bins the input vcfs into different categories. The comprehensive gold standard is used to evaluate metrics such as the sensitivity, precision and F-score for each bin. VISTA then uses a consensus-based approach, and decides the top performing caller for each bin (Supplemental tables 1-4). Next, VISTA merges the outputs of each of the top performing callers into one output vcf file. This approach ensures that the best caller per organism, variant type and variant length is selected, and the combination of several top callers exceeds existing callers. For VISTA’s discovery mode, where the ground truth is not known, we use the combination of callers identified during VISTA’s pretraining, on the organism closest to the input sample. We have found VISTA’s pre-trained algorithm to be generalizable across multiple samples of an organism.. If the user wants to compare the results of VISTA with other callers, and choses to provide a gold standard file, VISTA performs downstream analysis to produce statistics such as the sensitivity, precision and F1-score, as well as graphs comparing the output of VISTA to other variant callers.

### Compare deletion inferred from WGS data with the gold standard

We compared the deletions inferred from SV callers from WGS data (inferred deletions) with the molecular-based gold standard (true deletions). Start and end positions of the deletion were considered when comparing true deletions and inferred deletions. Inferred deletion was considered to be correctly predicted if the distance of right and left coordinates are within the resolution threshold τ from the coordinates of true deletion. We consider the following values for resolution threshold τ: 0 bp, 10 bp, 100 bp, 1000 bp,10000 bp. Since most tools had 0 matches when the threshold was kept at 0 bp, we starting threshold in the figures is kept to be 10bp. True positives were correctly predicted deletions, and were defined as deletions reported by the SV caller that were also present in the gold standard. In case an inferred deletion matches several true deletions, we randomly choose one of them. Similarly, in case true deletion matches several inferred deletions, we choose the first deletion that matches. The deletion predicted by the SV caller but not present in the golden standard was defined as false positives (FP). Similarly, each deletion present in the gold standard was matched with only one deletion predicted by the software. The SV that was not predicted by the SV caller were defined as false negatives (FN). SV detection accuracy was assessed using various detection thresholds (τ). The accuracy at threshold τ is defined as the percentage of SVs with an absolute error of deletion coordinates smaller or equal to τ. We have used the following measures to compare the accuracy of SV-callers:

● Sensitivity=TP/(TP+FN)
● Precision=TP/(TP+FP)
● F-score=2*Sensitivity*Precision/(Sensitivity+Precision)

### Compare computational performance of SV callers

The CPU time and RAM of each tool were measured to determine it’s computational performance. The statistics were measured for 1x coverage and full coverage bam files, with samples A/J and BALB/cJ for mouse data. The CPU time was computed using either the GNU time program that is inbuilt in make bash terminals or the Hoffman2 Cluster qsub command. For GNU time, we used this specific command /usr/bin/time -f “%e\t%U\t%S\t%M” which we either had to run manually on an interactive qsub session or through another method that wasn’t a qsub. This GNU time command would output one line containing Wallclock time in seconds, user-time in seconds, kernel-space time in seconds, and peak memory consumption of the process in kilobytes. CPU-time was calculated by adding user-time and kernel-space time. RAM usage was equivalent to peak memory consumption in the case of this command. For qsubs on the Hoffman2 Cluster, we used the command qsub -m e which would email the user a full list of records when the tool finished running. This list included CPU-time and Max mem which was designated as RAM usage for each tool.

### Downsample the WGS samples

We have used a custom script to downsample the full coverage BAM file to desired coverage. Existing tools (e.g., samtools) are not suitable for this purpose as they treat each read from a read pair independently, resulting in singletons reads in the downsample BAM file.

### VISTA Train-test experiments

For deletions, we trained VISTA on 5 different mouse strains on chr19 (A_J, AKR_J, BALB_CJ, C3H_HeJ, CBA_J) to find best individual callers per length bin, and tested on DBA_2J and LP_J. For human data, we trained VISTA on chr 17, chr 18, chr 19, and tested on chr 15 and chr16.

## Data availability

WGS mouse strains for the samples A/J, AKR/J, BALB/cJ, CBA/J, C3H/HeJ, DBA/2J and LP/J used for benchmarking of SV-callers are available under the following accession numbers in the European Nucleotide Archive: ERP000038, ERP000037, ERP000039, ERP000040, ERP000044, and ERP000045. The output VCF’s produced by the tools, the gold standard VCF’s, the analysis scripts, figures, and log files are available at https://github.com/Mangul-Lab-USC/benchmarking_SV. The human high confidence bed file can be found here https://www.nist.gov/programs-projects/genome-bottle

The novoaligned bams data for the HG002_NA24385 son genome were downloaded from https://ftp-trace.ncbi.nlm.nih.gov/ReferenceSamples/giab/data/AshkenazimTrio/HG002_NA24385_son/NIST_Illumina_2×250bps/novoalign_bams/

The 10 human sample (HG00733, HG00438, HG00621, HG00735, HG00741, HG01071, HG01106, HG01109, HG01243, HG01175) bams data are from the 1000 Genomes Project with 30x coverage on the reference GRCh38^39^. They can be downloaded through The International Genome Sample Resource (IGSR)^42^ portal at https://www.internationalgenome.org/data-portal/sample.

## Code availability

All code required to produce the figures and analysis performed in this paper is freely available at https://github.com/Mangul-Lab-USC/VISTA_paper

## Tool Availability

The source code for VISTA is open source under the MIT license and is available at: https://github.com/Mangul-Lab-USC/VISTA

## Supplementary Materials

### Supplementary Tables

**Table S1:** Train-test distribution for human HG002 WGS data. VISTA was trained on chromosomes 17-19 to determine the highest performing caller per length bin and was tested on chromosomes 15 and 16.

| Strain | 50-100 | 100-500 | 500-1000 | 1000+ | Results [VISTA, Parliament2, Survivor] |
| --- | --- | --- | --- | --- | --- |
| CHR17 | Octopus | Manta | Delly | Delly | [0.77,0.21,0.53] |
| CHR18 | Octopus | Manta | Delly | GENOMESTRIP | [0.89,0.45,0.71] |
| CHR19 | Octopus | Manta | Delly | GENOMESTRIP | [0.71,0.21,0.46] |
| **CHR15** | **Octopus** | **Manta** | **Delly** | **GENOMESTRIP** | **[0.75,0.44,0.71]** |
| **CHR16** | **Octopus** | **Manta** | **Delly** | **GENOMESTRIP** | **[0.73,0.23,0.67]** |

**Table S2:**
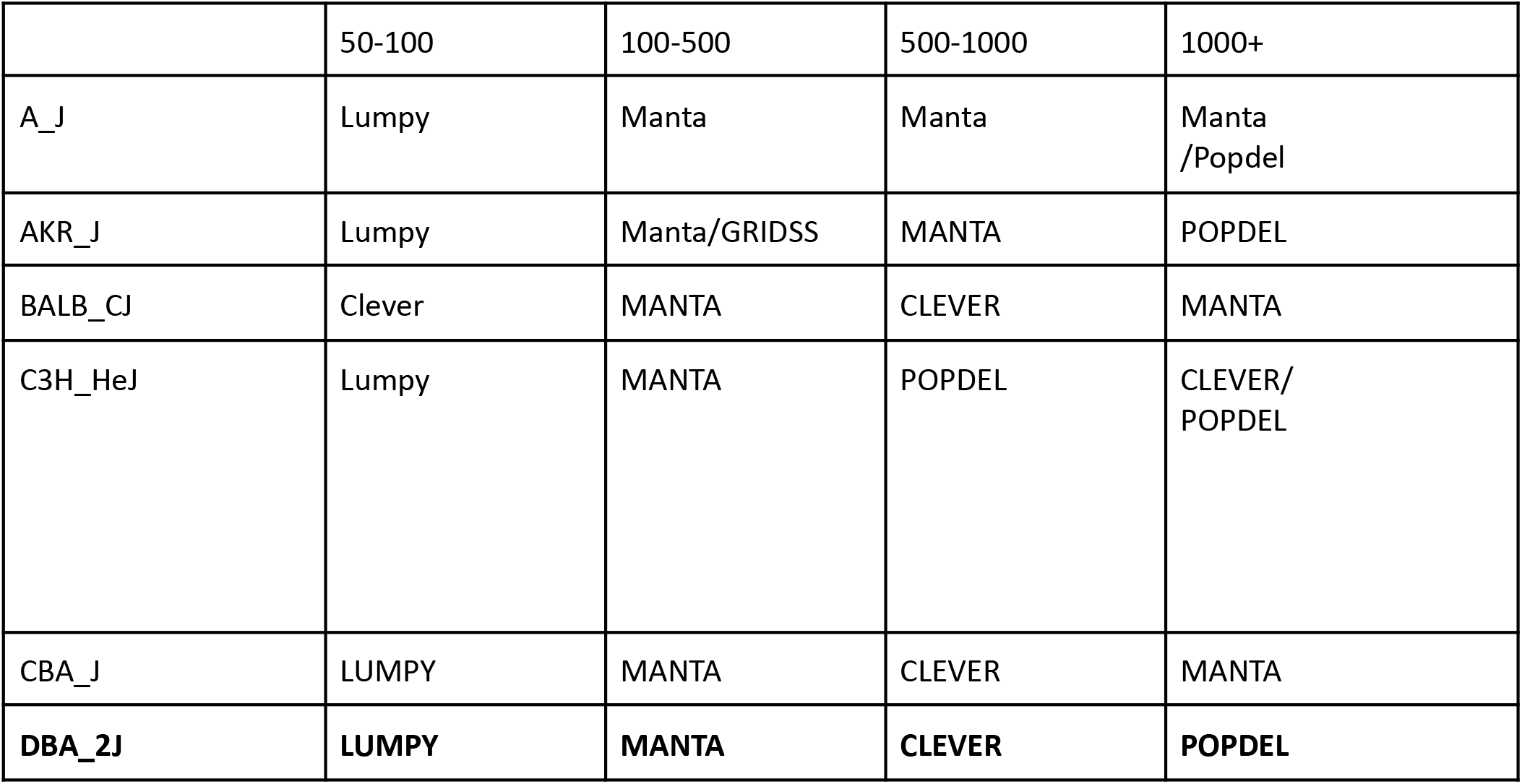
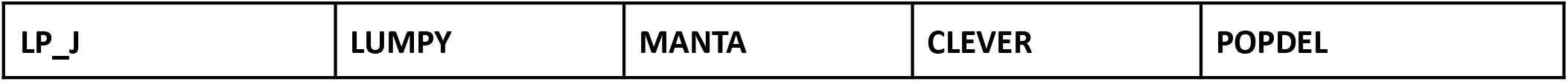
Train-test distribution for 7MM. VISTA was trained on 6 different mouse strains to determine the highest performing caller per bins and was tested on chromosomes DBA_2J and LP_J.

|  | 50-100 | 100-500 | 500-1000 | 1000+ |
| --- | --- | --- | --- | --- |
| A_J | Lumpy | Manta | Manta | Manta /Popdel |
| AKR_J | Lumpy | Manta/GRIDSS | MANTA | POPDEL |
| BALB_CJ | Clever | MANTA | CLEVER | MANTA |
| C3H_HeJ | Lumpy | MANTA | POPDEL | CLEVER/ POPDEL |
| CBA_J | LUMPY | MANTA | CLEVER | MANTA |
| **DBA_2J** | **LUMPY** | **MANTA** | **CLEVER** | **POPDEL** |

**Table S3:**
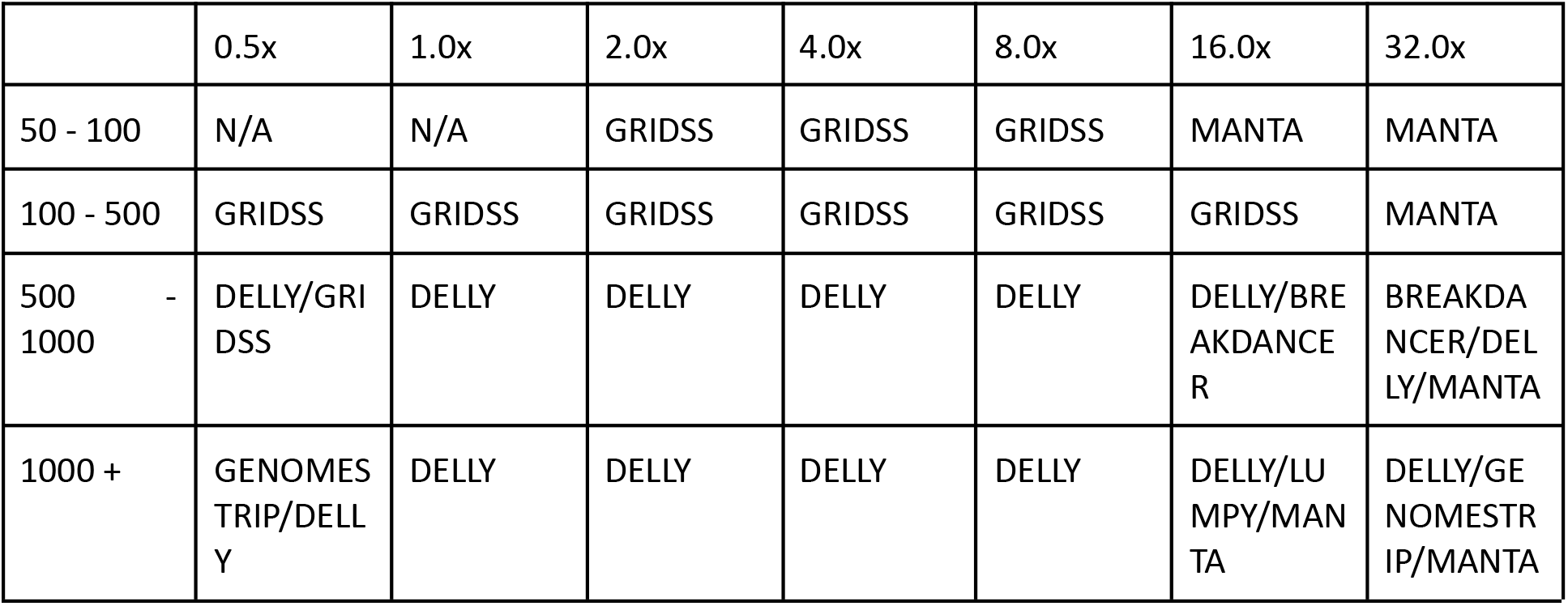
Training based on the f-score for downsampled data.

|  | 0.5x | 1.0x | 2.0x | 4.0x | 8.0x | 16.0x | 32.0x |
| --- | --- | --- | --- | --- | --- | --- | --- |
| 50 - 100 | N/A | N/A | GRIDSS | GRIDSS | GRIDSS | MANTA | MANTA |
| 100 - 500 | GRIDSS | GRIDSS | GRIDSS | GRIDSS | GRIDSS | GRIDSS | MANTA |
| 500 - 1000 | DELLY/GRIDSS | DELLY | DELLY | DELLY | DELLY | DELLY/BREAKDANCER | BREAKDANCER/DELLY/MANTA |
| 1000 + | GENOMES TRIP/DELLY | DELLY | DELLY | DELLY | DELLY | DELLY/LUMPY/MANTA | DELLY/GENOMESTRIP/MANTA |

**Table S4:** Train-test distribution for HPRC-HC. VISTA was trained on HG00438, HG00621, HG00733 to determine the highest performing caller per length bin.

|  | 50-100 | 100-500 | 500-1000 | 1000+ |
| --- | --- | --- | --- | --- |
| HG00438 | Octopus | Manta | DELLY | Manta/Smoove |
| HG00621 | Octopus | Manta | DELLY | Manta/Smoove |
| HG00733 | Octopus | Manta | DELLY | Manta/Smoove |

**Table S5:** Gold standard deletion calls from chromosome 19 from 7 inbred mouse strains.

| Strain name | Total number<br>of gold<br>standard<br>deletions | Average<br>deletion length | Minimum<br>deletion length | Maximum<br>deletion length |
| --- | --- | --- | --- | --- |
| A/J | 533 | 1968 | 15 | 239572 |
| AKR/J | 504 | 1400 | 13 | 24301 |
| BALB/cJ | 545 | 1947 | 15 | 239572 |
| C3H/HeJ | 539 | 1993 | 15 | 239572 |
| DBA/2J | 609 | 1301 | 16 | 24301 |
| LP/J | 483 | 1534 | 13 | 24301 |
| CBA/J | 586 | 1330 | 24 | 24301 |

**Table S6:** Parliament2 results on different human samples. Parliament2 outputs results including different callers.

| Samples | Breakdancer | CNVnator | Delly2 | Lumpy | Manta | BreakSeq2 |
| --- | --- | --- | --- | --- | --- | --- |
| HG00733 | Yes | Yes | Yes | Yes | No | No |
| HG00438 | Yes | Yes | Yes | Yes | No | No |
| HG00621 | Yes | Yes | Yes | Yes | Yes | No |
| HG00735 | No | Yes | Yes | Yes | Yes | Yes |
| HG00741 | No | Yes | Yes | Yes | Yes | Yes |
| HG01071 | Yes | Yes | Yes | Yes | No | No |
| HG01106 | Yes | Yes | Yes | Yes | No | No |
| HG01109 | Yes | Yes | Yes | Yes | Yes | No |
| HG01243 | No | Yes | Yes | Yes | Yes | Yes |
| HG01175 | No | Yes | Yes | Yes | Yes | Yes |
| HG002 | No | Yes | Yes | Yes | Yes | No |

### Supplementary Figures

**Supplementary Figure 1:**
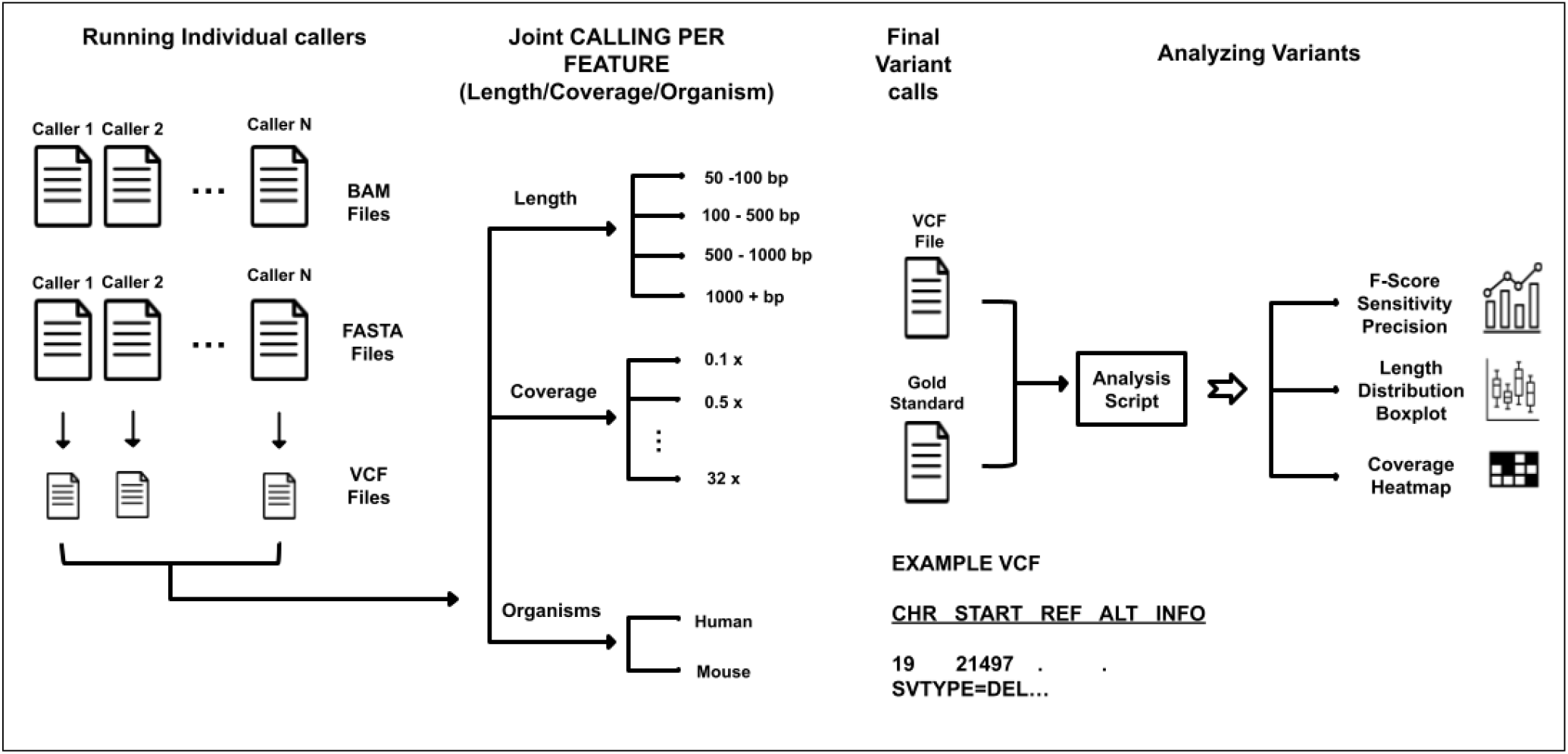
Overview of the approach implemented in VISTA.

**Supplementary Figure 2:**
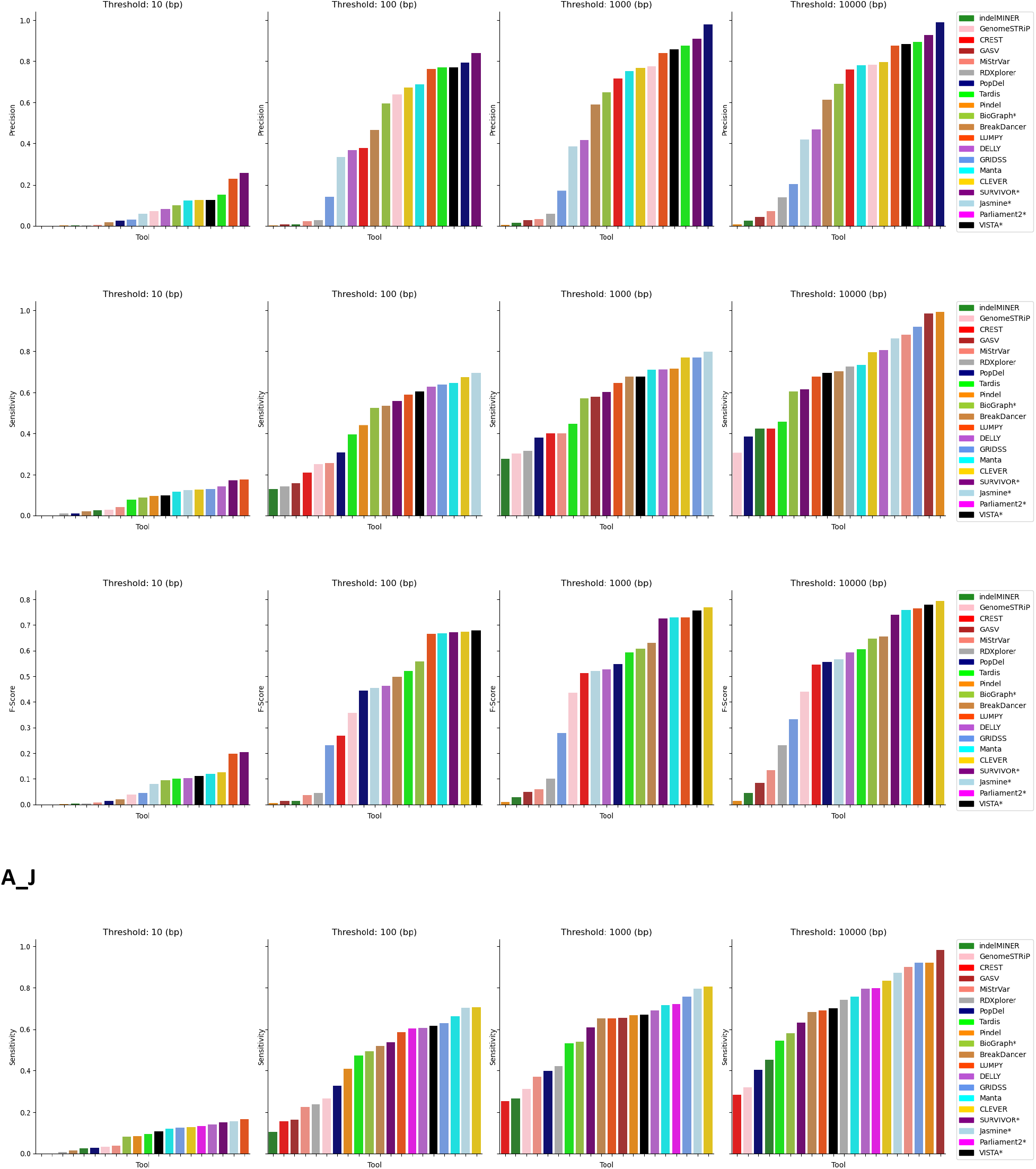
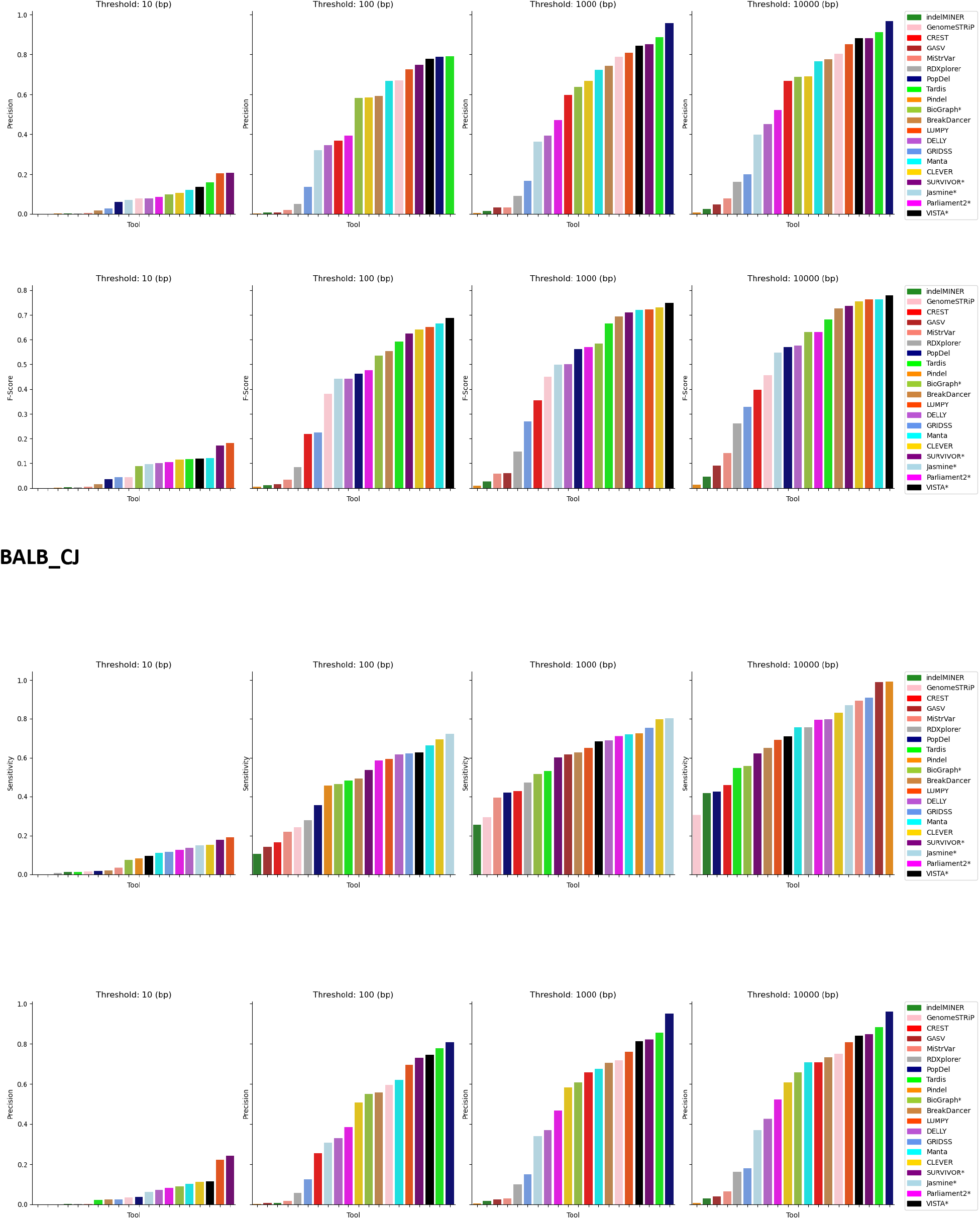
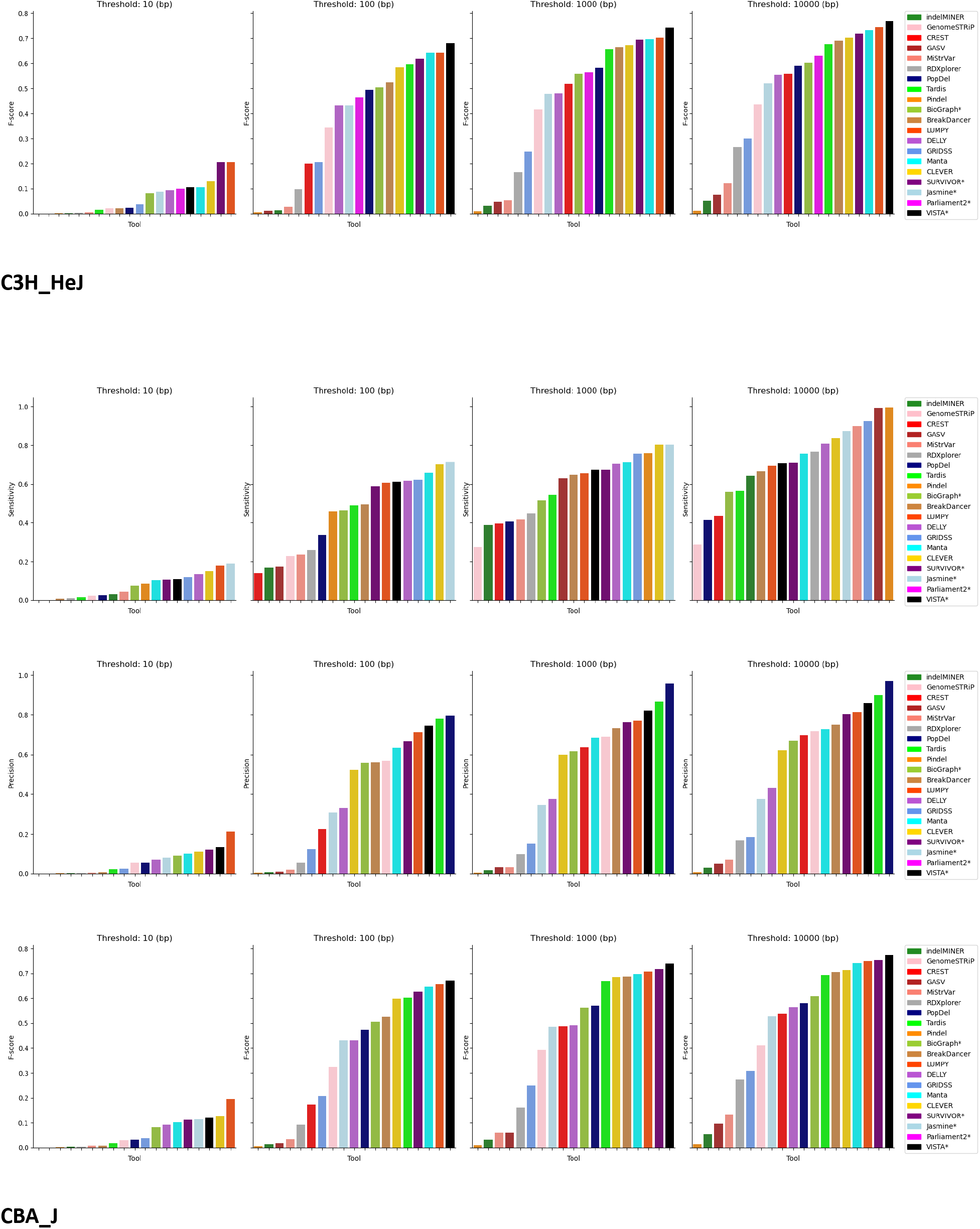
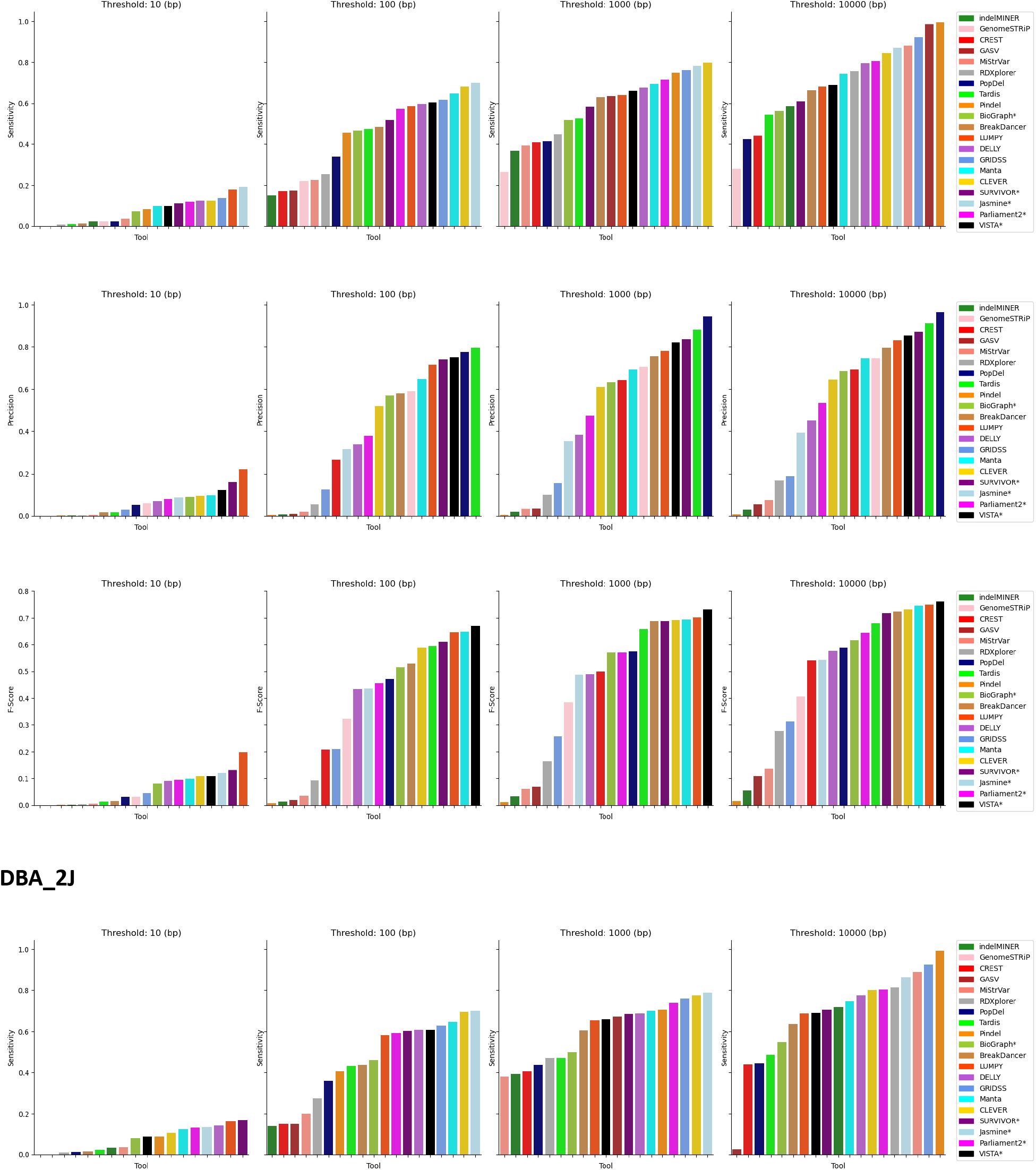
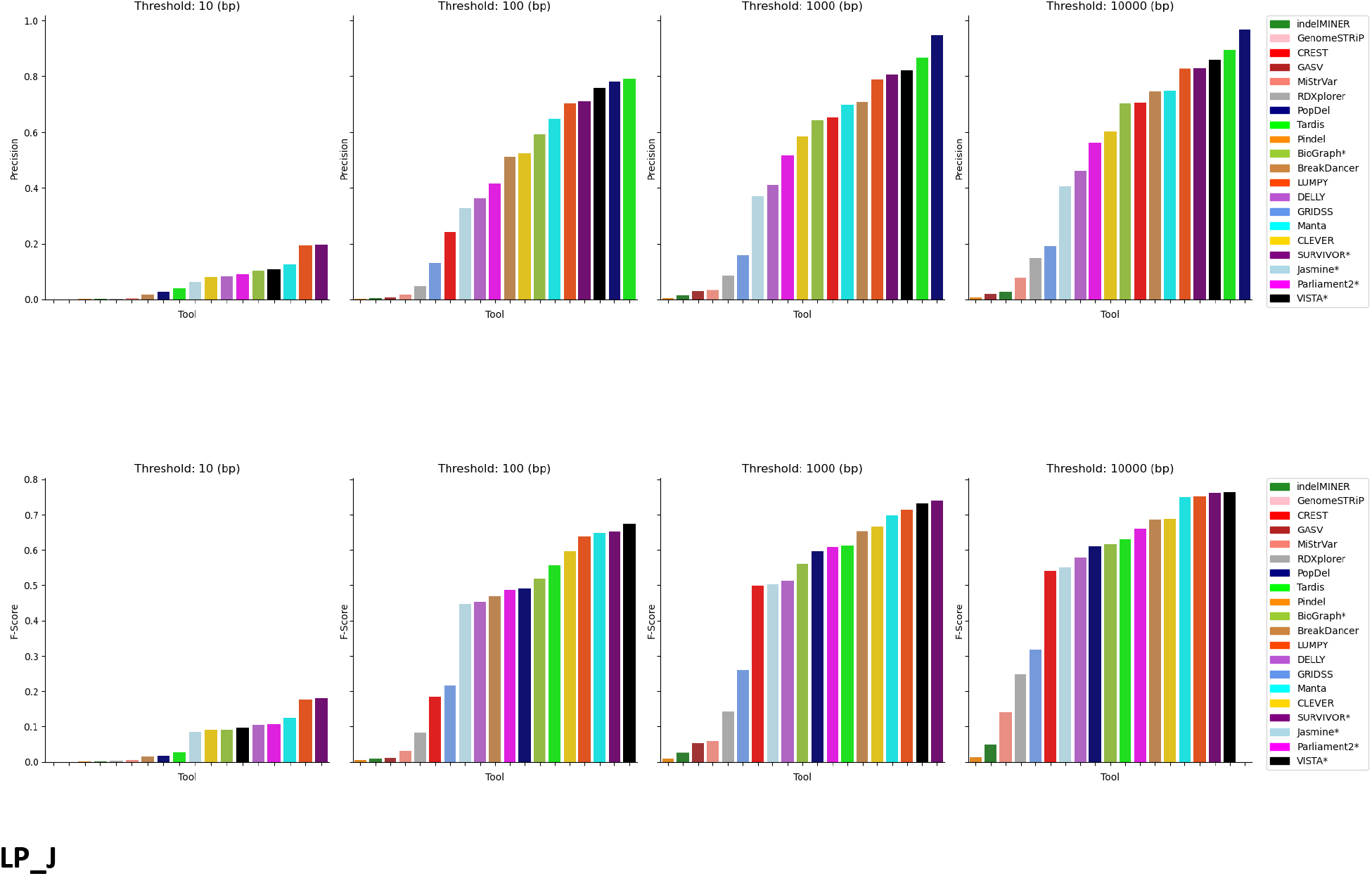
Comparison of the performance of VISTA vs other callers for the 7 individual mouse strains (7MM)

**Supplementary Figure 3:**
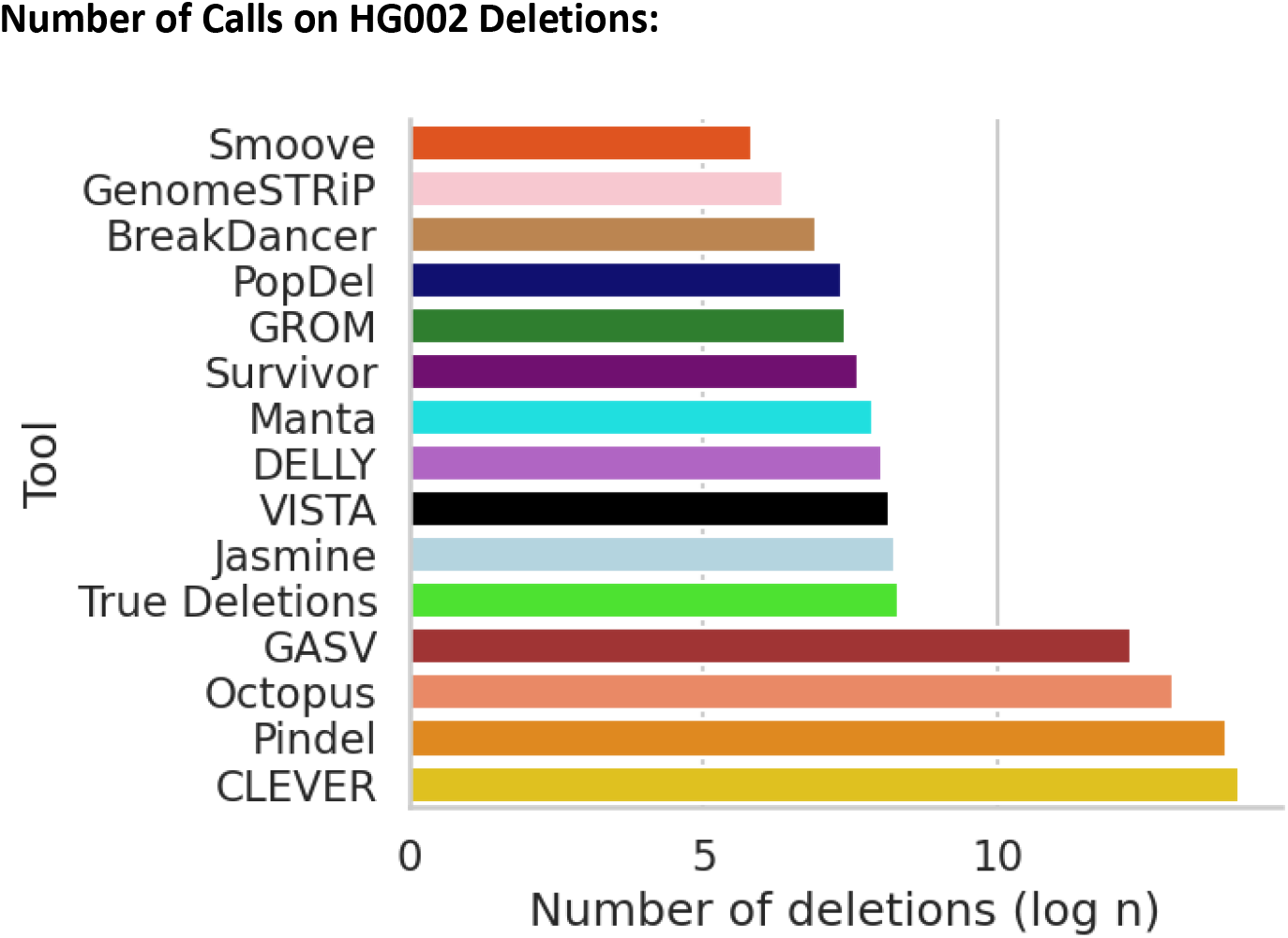
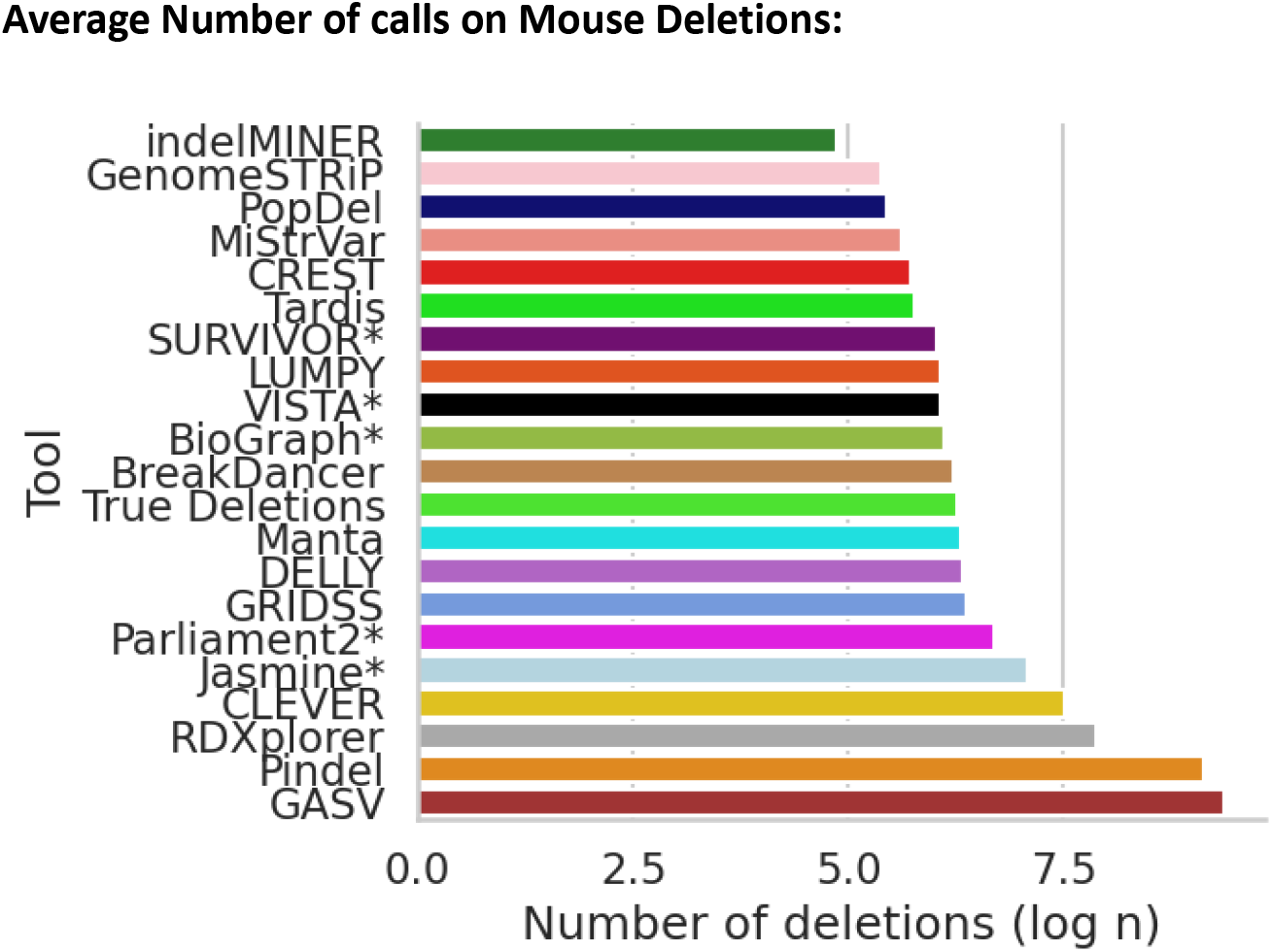
Number of molecularly-confirmed deletions by VISTA (black), ground truth (green) and other SV callers for a) GIAB-HC dataset and b)MM-7 dataset.

**Supplemental Figure 4:**
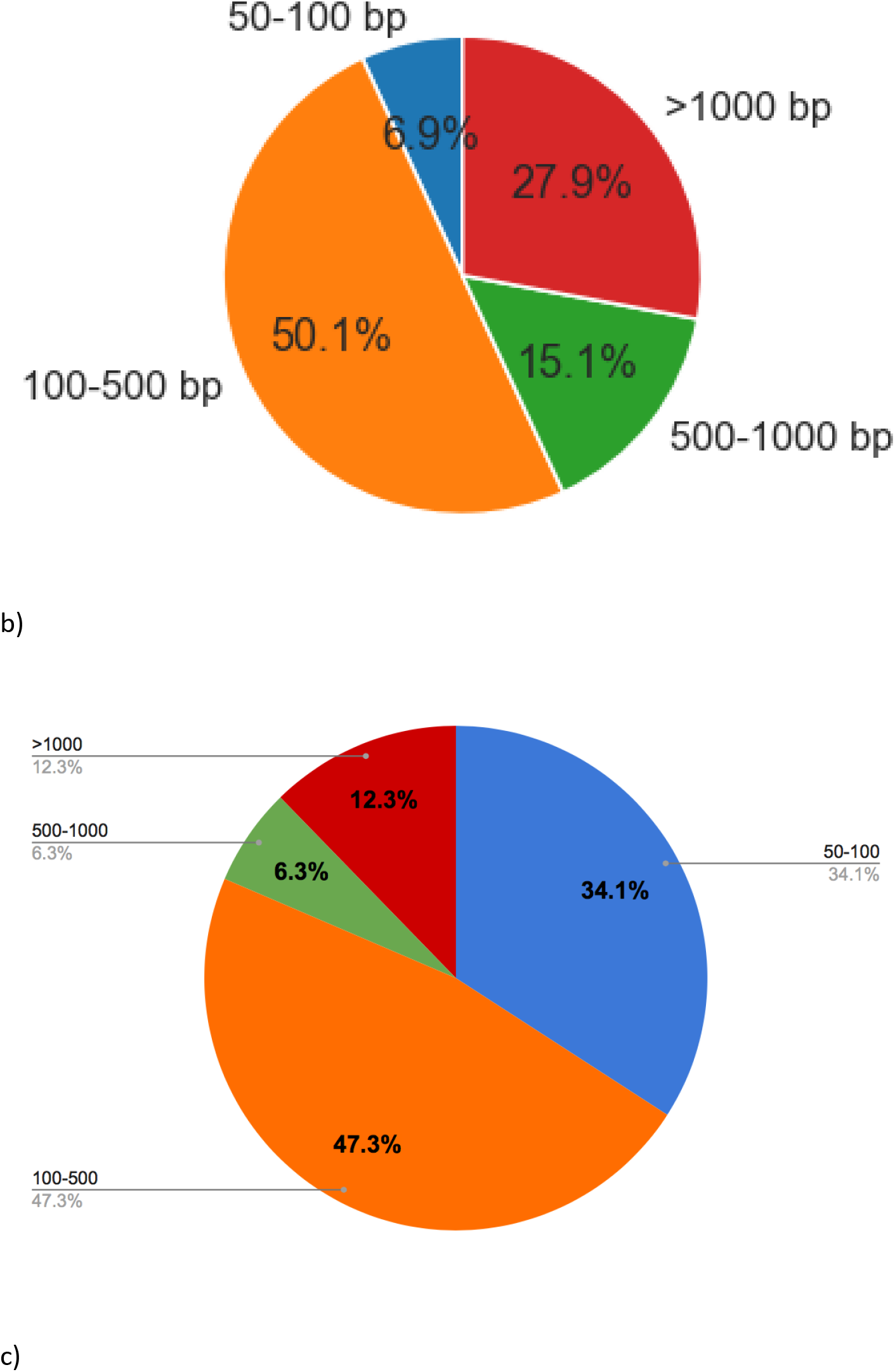
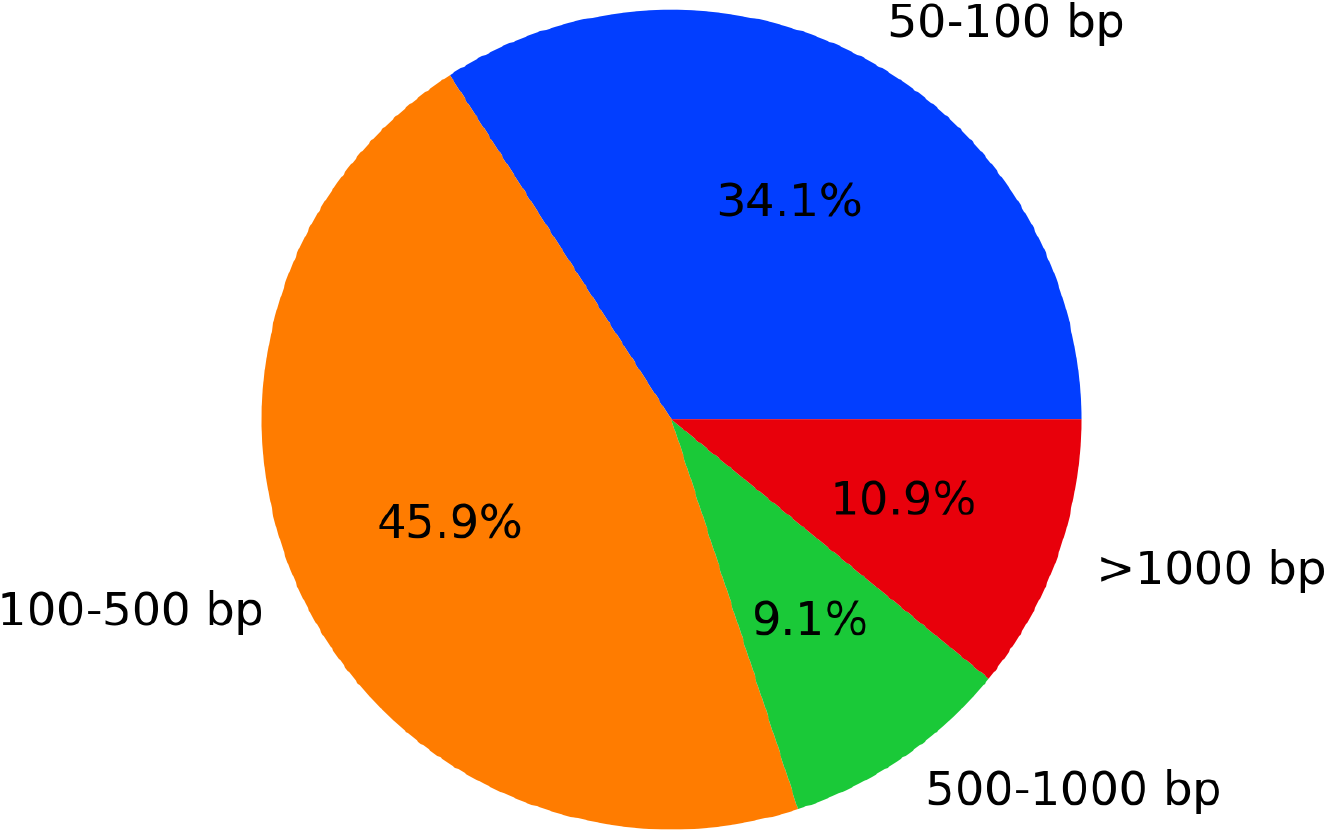
Total number of deletions across a) MM-7 b) GIAB-HC c)HPRC-HC.

**Supplementary Figure 5:**
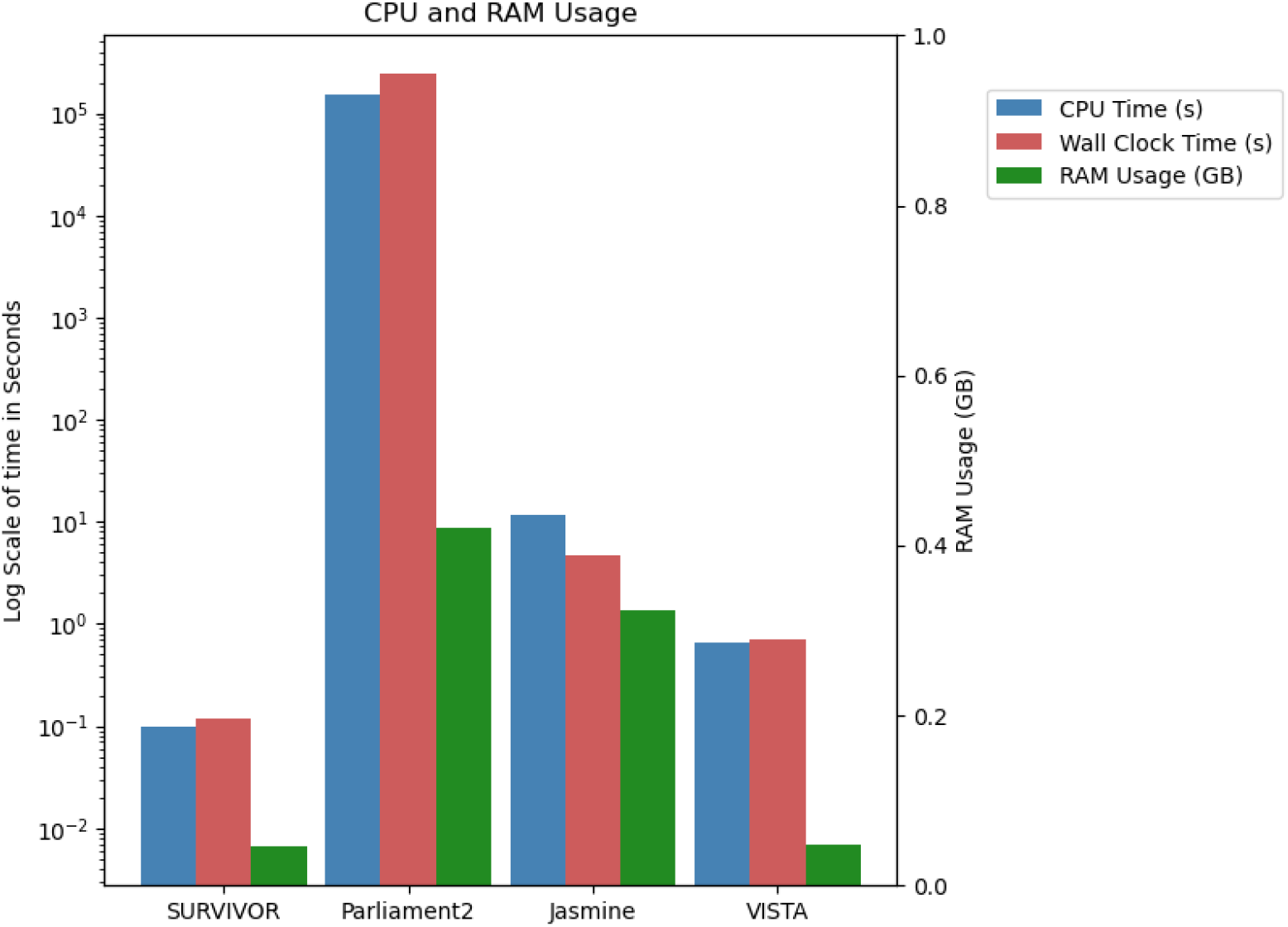
A comparison of the computational performance of SV consensus-based callers. The bar plot depicts the CPU and RAM usage across all of the tools.

**Supplementary Figure 6:**
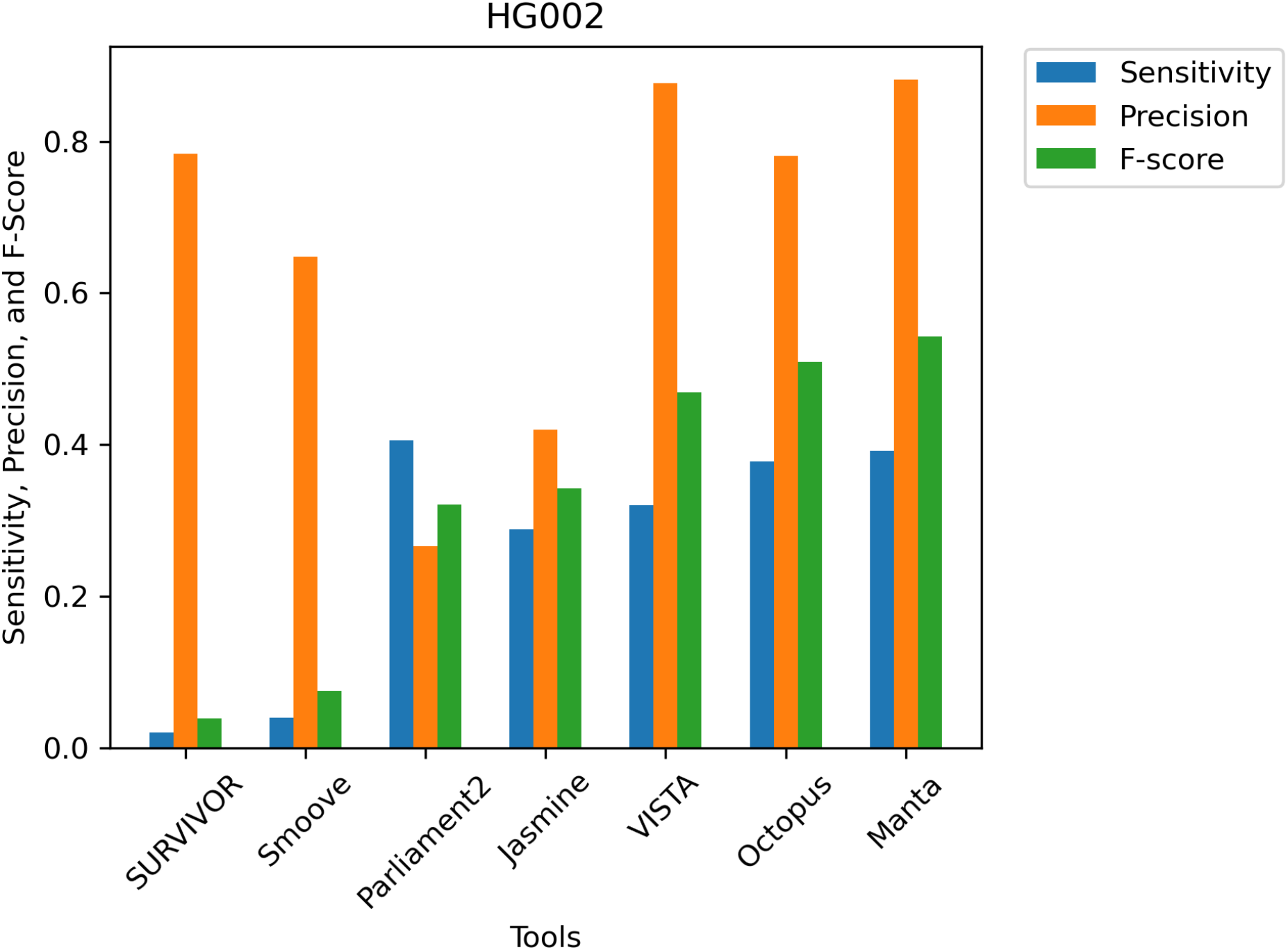
Comparing the performance of VISTA pre-trained on HG002 on HGDP data compared to other individual and consensus callers at 100 bp threshold.

